# Comparative immune responses to *Mycobacterium tuberculosis* in people with latent infection or sterilizing protection

**DOI:** 10.1101/2022.06.13.495865

**Authors:** Emilie Jalbert, Cuining Liu, Vidya Mave, Nancy Lang, Anju Kagal, Chhaya Valvi, Mandar Paradkar, Nikhil Gupte, Rahul Lokhande, Renu Bharadwaj, Vandana Kulkarni, Amita Gupta, Adriana Weinberg

## Abstract

Tuberculosis (TB) affects 2 billion people worldwide and causes 1.5 million deaths every year. There is great need for new TB vaccines that are more efficacious than the currently licensed BCG vaccine, which provides only limited protection. Our goal was to identify potential targets for new TB vaccines by characterizing the immune responses that distinguish individuals with sterilizing protection against TB (TB-resisters), defined by presence of TB-specific immune responses and absence of latent infection, from individuals with latent TB infection (LTBI-participants).

Cryopreserved peripheral blood mononuclear cells (PBMC) from 13 TB-resisters and 10 LTBI-participants were analyzed by high dimensional spectral flow cytometry after overnight *M. tuberculosis (Mtb)* antigenic stimulation or unstimulated control. Activation of conventional and nonconventional T cells, NK cells, and antigen presenting cells (APC) was compared between the two groups. Compared with LTBI-participants, TB-resisters had significantly higher proportions of conventional and nonconventional T cells expressing granzyme B (GranzB) and PD-1, and of polyfunctional cells in unstimulated and *Mtb*-stimulated conditions. Conversely LTBI-participants had higher expression of CD25, CD69, CD107a, IL10 and IFNγ. An unbiased cluster analysis revealed higher frequency of recently described CD8+GMM+GranzB+ T cells in unstimulated PBMC from TB-resisters than LTBI-participants. APC activation revealed very few differences between TB-resisters and LTBI-participants. An exploratory analysis of responses in 14 BCG-recipients with minimal exposure to TB showed multiple differences with TB-resisters and LTBI-participants in PBMC activation; lower polyfunctionality of T cells and APC in *Mtb*-stimulated PBMC; and absence of CD8+GMM+GranzB+ T cells.

We conclude that combined increased T cell expression of GranzB and checkpoint inhibitors may contribute to immune protection against TB and may be targeted by new vaccines.

## Introduction

Tuberculosis (TB) is a major global health problem, since approximately 2 billion people are infected with *M. tuberculosis (Mtb).* Vaccines are the most powerful tools for limiting the morbidity and mortality of many infectious diseases, but Bacillus Calmette-Guérin (BCG)—the only licensed TB vaccine—confers limited protection^1–7^. BCG administered to infants at birth is ~80% effective in preventing disseminated and central nervous system TB. In contrast, BCG has variable efficacy against pulmonary TB when administered to infants or adults, and many studies could not demonstrate any efficacy^1–8^. BCG generates TB-specific Th1 cell-mediated immunity (CMI) that has been considered essential for protection against TB, although this concept has been disputed by recent studies^9–11^. For example, recent vaccine candidates that generate more robust *Mtb*-specific Th1 CMI than BCG did not have greater efficacy against pulmonary TB than BCG^9,12–14^. Additional evidence that *Mtb* Th1 responses may not predict protection against *Mtb* infection is that 10 to 20% of the individuals with latent TB infection, as defined by the presence of *Mtb* CMI measured by tuberculin skin test (TST) or IFNγ release assays (IGRA), a measure of Th1 immunity, develop active TB disease over time. In contrast, 20% of individuals with household contacts with highly contagious active pulmonary TB infection do not develop clinical disease and have negative TST or IGRA^15^. These individuals, designated as TB-resisters, have antibodies and limited CMI responses to *Mtb,* confirming prior *Mtb* infection, but their T cells do not produce IFNγ in response to *Mtb* ex vivo restimulation^16,17^. They also have genetic traits that differentiate them from people with LTBI ^18–21^. TB-resisters are deemed to have sterilizing immunity against *Mtb*^22,23^. However, the mechanisms of protection against *Mtb* infection in TB-resisters are incompletely understood^17,24^. Identifying these mechanisms would benefit the development of new vaccines that may confer sterilizing immune protection against *Mtb*^25^.

In addition to adaptive CMI, innate immunity is a critical mechanism of protection against infections. Innate immunity is rapidly deployed and constitutes the first line of immune-mediated defense. Moreover, innate immune responses critically contribute to the development of antigen-specific adaptive CMI and establish a feedback mechanism that allows them to be also boosted by adaptive CMI^26–32^. Persistent memory-like innate immune responses have been described against viruses, tumors, and other antigens, including *Mtb*^28,32–43^.

Natural killer (NK) cells, γδ T cells, NKT cells, invariant NKT (iNKT) cells, macrophages, monocytes (mono) and dendritic cells (DC) have been shown to undergo clonal expansions and/or epigenetic modifications after exposure to immunogens that can confer antigen specificity and/or allow them to activate transcription programs that improve their functionality^43,44^. Recent studies described a germline encoded, mycolyl-reactive iNKT cell subset (GEMT) that recognizes the lipid glucose monomycolate in the context of CD1b and participates in the elimination of *Mtb* from the host^40,42,45^. Compared to other iNKT, CD1b-restricted GEMT cells have a more diverse T-cell receptor repertoire, which permitted the identification of GEMT cell clonal expansions in patients recovering from TB and their persistence in the host for several years after *Mtb* elimination^40,42^. Another nonconventional T cell subset of interest is the mucosal-associated invariant T cells (MAIT) that developmentally share some characteristics both with iNKT and γδ T cells. MAIT can recognize microbial riboflavin derivatives presented by the major histocompatibility-related receptor 1 (MR1) and play a critical role in antibacterial, including *Mtb,* pulmonary defenses via cytokine production and cytotoxicity^46–48^. MR1+ MAIT were shown to account for most IFNγ production in response to BCG^49^. MR1-MAIT lack the receptor for antigen recognition and are deemed to respond to secreted cytokines^50,51^. The role of innate immune responses in sterilizing immunity against *Mtb* infection has not been studied.

The overarching goal of our study was to identify innate and adaptive immune responses that differentiate TB-resisters from people with LTBI (LTBI-participants) among household contacts of active pulmonary TB cases. We leveraged a longitudinal cohort study of household contacts of individuals with active pulmonary TB disease [Cohort For TB Research By The Indo-US Medical Partnership Multicentric Prospective Observational Study (C-TRIUMPH)] in Pune, India, by using advanced spectral flow cytometry technology and highdimensional analytic tools on peripheral blood mononuclear cells (PBMC) archived from the study participants^23,52,53^. To further understand how the responses to *Mtb* in LTBI-participants and TB-resisters may differ from those induced by BCG, we also included in our analysis a group with documented BCG vaccination and with minimal exposure to TB.

## Results

### Characteristics of the study population

The study used PBMC from 13 TB-resisters and 11 LTBI-participants in C-TRIUMPH, who had intense exposure to active pulmonary TB household contacts (**Table 1**). Importantly, TB-resisters and LTBI-participants had similarly high exposure scores to TB (median=7), a measure of the likelihood of acquiring *Mtb* infection from a household index case^54^. Also recruited were 14 BCG-recipients with limited exposure to TB by virtue of having spent most of their lives in the United States or other countries with low incidence of TB. In addition to these differences in upbringing and environmental exposures, we noted that BCG-recipients significantly differed from C-TRIUMPH participants in race/ethnicity, but not in age or BMI.

**Table 1.** Characteristics of the study population.

|  | TB-Resisters | LTBI+<br>participants | BCG | P-Value*<br>(TB-Resisters<br>vs. LTBI+) | P-Value*<br>(three-<br>group) |
| --- | --- | --- | --- | --- | --- |
| N | 13 | 11 | 14 |  |  |
| Median years of age (range) | 24 (9, 65) | 29 (7, 62) | 29 (20, 41) | 0.81 | 0.71 |
| N Female sex (%) | 6 (46) | 4 (36) | 10 (71.4) | 0.70 | 0.19 |
| N Race and Ethnicity (%) |  |  |  |  |  |
| Asian (Indian subcontinent) | 13 (100) | 11 (100) | 0 (0) |  |  |
| Asian (not Indian subcontinent) | 0 (0) | 0 (0) | 3 (21.4) |  |  |
| Black | 0 (0) | 0 (0) | 2 (14.3) | na | <0.0001 |
| Hispanic Non-Black | 0 (0) | 0 (0) | 5 (35.7) |  |  |
| Non-Hispanic White | 0 (0) | 0 (0) | 4 (28.6) |  |  |
| Median BMI<br>(range) | 23.3<br>(13.1, 30.5) | 20.4<br>(13.5, 29.8) | 22.8<br>(20, 30.6) | 0.49 | 0.49 |
| Median exposure score (range) | 7 (4, 9) | 7 (4, 9) | na | 0.84 | na |
| N with BCG scar (%) | 9 (64.3) | 5 (35.7) | na | 0.24 | na |

\* ANOVA for continuous variables and Fisher's exact for categorical variables.
**Abbreviations:** BCG=Bacille de Calmette-Guerin; BMI=Body mass index (kg/m<sup>2</sup>); LTBI=latent tuberculosis infection; N=number; na=not applicable

### Phenotypic and functional characteristics of innate and adaptive immune responses in TB-resisters and LTBI-participants

We used two high dimensional flow cytometry panels [T-cell and antigen-presenting-cell (APC)] on PBMC stimulated with *Mtb* antigen or unstimulated controls (**Table S1**). In the <u>T-cell panel</u>, we characterized CD4+ and CD8+ conventional T cells (Tconv), NKT, γδ T, GEMT, iNKT, MR1+ and MR1-MAIT, and NK cells. Functionality was assessed by expression of CD25 and CD69 activation markers; PD1 immunologic checkpoint receptor; CD107a (lysosome degranulation) and granzyme B (GranzB) cytotoxicity markers; GMCSF, IL2, IL10, IL17, IFNγ, and TNFα cytokines; and Ki67 proliferation marker. In the <u>APC panel</u>, we characterized functionality of CD14+ monocytes (Mono), CD123+ plasmacytoid DC (pDC), CD11c+CD14-total conventional DC (cDC total), CD141+ cDC1 and CD1c+ cDC2 subsets using the following functional readouts: CD40, CD80 and CD83 activation markers; PDL1 immunologic checkpoint ligand; and GMCSF, IL1β, IL8, IL10, IL12p40, IL27 and TNFα cytokines. Gating trees are shown in **Fig S1**.

In *unstimulated* conditions, 10 functional cell subsets had significantly higher frequencies in TB-resisters compared with LTBI-participants, consisting of GranzB-expressing CD4+ Tconv, CD8+ Tconv, CD8+ iNKT and CD8+ NKT; PD1-expressing CD4+ Tconv, CD8+ Tconv, CD8+ iNKT and MR1-MAIT; CD40+ pDC; and IL1β+ Monos (**Fig 1**). In contrast, we only detected a single subset (Ki67+MR1+ MAIT) with significantly higher frequencies in LTBI-participants compared with TB-resisters.

**Figure 1.**
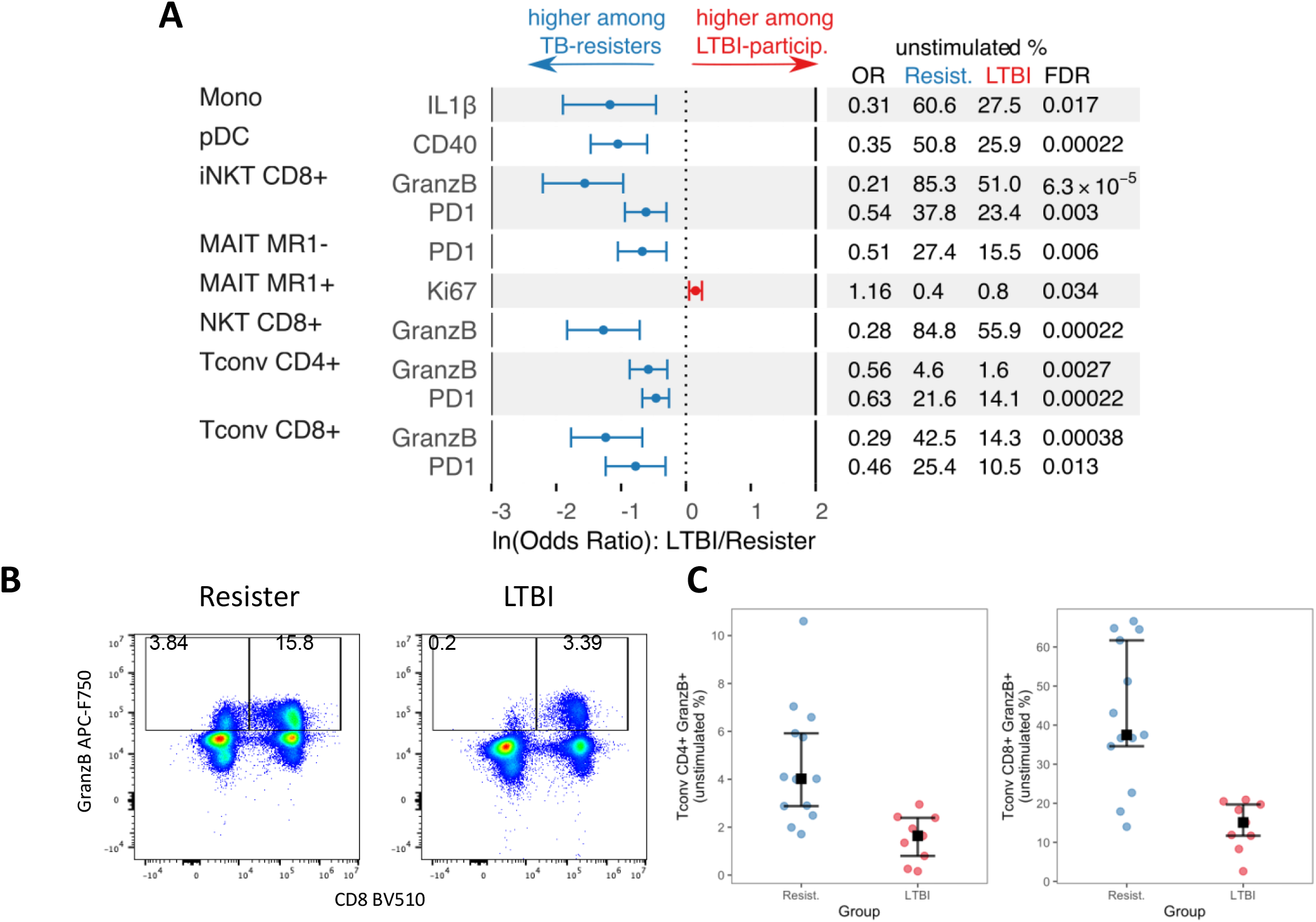
Functional phenotypes differentially expressed in unstimulated PBMC from TB-resisters and LTBI-participants. Data were derived from 13 TB-resisters and 11 LTBI-participants. PBMC were analyzed after overnight rest in culture medium. **A**. Forest plot displays ln-transformed odds ratios (lnOR) and 95% confidence intervals (CI). The dotted line shows no average effect (lnOR = 0, corresponding to OR = 1). Features on the right side of this line are more highly expressed among LTBI-participants (OR > 1), and features on the left side are more highly expressed among TB-resisters (OR < 1). The table shows the absolute OR and the means of each parameter in TB-resisters and LTBI-participants. **B.** Typical scatter plots exemplifying differences in GranzB+ CD4+ and CD8+ Tconv between TB-resisters (blue dots) and LTBI-participants (red dots). **C**. Dot plots showing frequencies of GranzB+ CD4+ and CD8+ Tconv in all participants.

Many of the functional differences between TB-resisters and LTBI-participants observed in unstimulated PBMC persisted after *Mtb* ex vivo stimulation (**Fig 2**). Differences between TB-resisters and LTBI-participants in *Mtb-stimulated* PBMC included higher proportions of (1) GranzB-expressing CD4+ Tconv, CD8+ Tconv, CD8+ iNKT. CD8+ NKT and MR1-MAIT cells; and (2) PD1-expressing CD4+ Tconv, CD8+ Tconv and MR1-MAIT in TB-resisters; and higher proportions of (1) CD25-expressing CD4+ and CD8+ Tconv and NKT, CD8+ iNKT, and MAIT MR1-; (2) CD107+CD8+ iNKT and NKT; (3) IFNγ-expressing CD4+ and CD8+ iNKT; (4) CD8+IL10+ iNKT; and (5) CD69+ γδ T cells in LTBI-participants.

**Figure 2.**
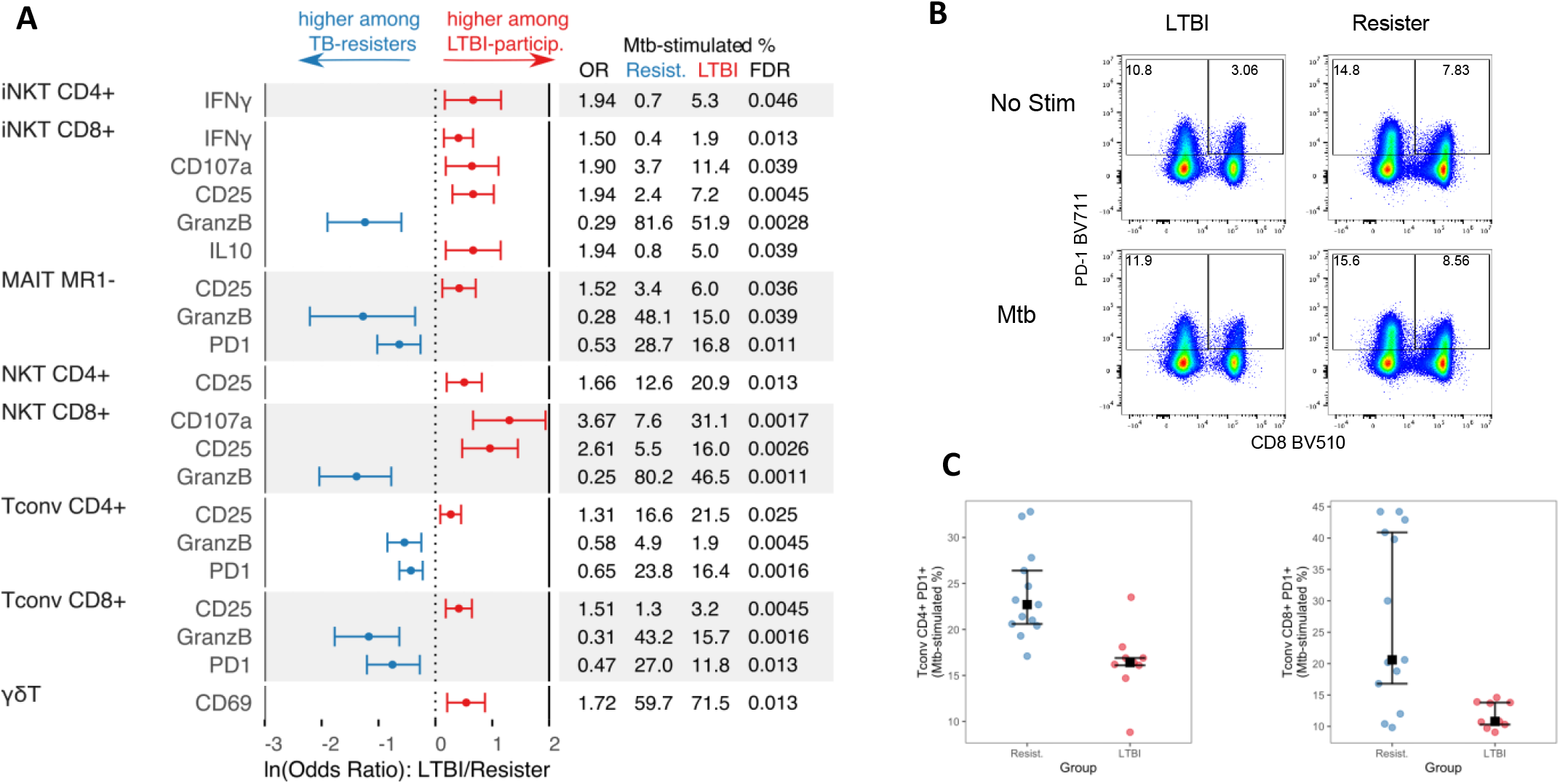
Functional phenotypes differentially expressed in *Mtb*-stimulated PBMC from TB-resisters and LTBI-participants. Data were derived from 13 TB-resisters and 9 LTBI-participants that contributed PBMC to the T cell analysis. PBMC were analyzed after pre-optimized overnight stimulation with *Mtb* cell membrane (see methods). **A**. Forest plot displays natural log (ln)-transformed odds ratios (lnOR) and 95% confidence intervals (CI). The dotted line shows no average effect (lnOR = 0, corresponding to OR = 1). Features on the right side of this line are more highly expressed among LTBI-participants (OR > 1), and features on the left side are more highly expressed among TB-resisters (OR < 1). The table shows the absolute OR and the means of each parameter in TB-resisters and LTBI-participants. **B.** Typical scatter plots exemplifying differences in PD1+ CD4+ and CD8+ Tconv between TB-resisters and LTBI-participants. **C**. Frequencies of GranzB+ CD4+ and CD8+ Tconv in all participants, including medians, upper and lower quartiles marked on the graph.

*Mtb-memory* responses were characterized by subtracting frequencies of unstimulated from *Mtb-stimulated* corresponding PBMC. Using this definition, we identified 6 differentially expressed functional subsets with significantly higher frequencies in LTBI-participants compared with TB-resisters: CD25+ and CD69+ γδ T cells; CD4+CD25+ Tconv; CD8+IFNγ+ iNKT; and CD8+CD107a+ NKT (**Fig 3**). There were no *Mtb-memory* subsets with higher frequency in TB-resisters than in LTBI-participants.

**Figure 3.**
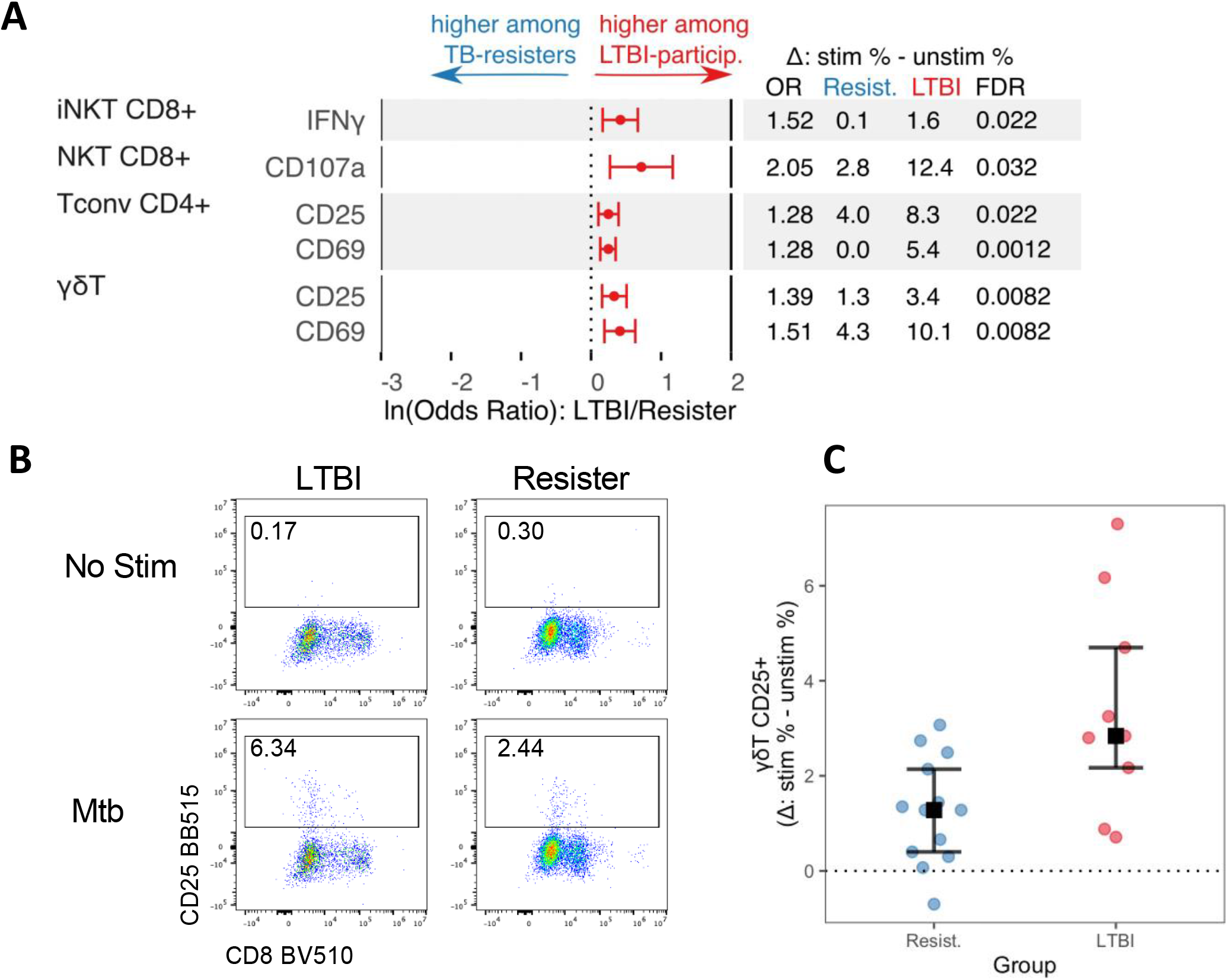
*Mtb*-specific memory responses in TB-resisters and LTBI-participants. Data were derived from 13 TB-resisters and 9 LTBI-participants that contributed PBMC to the T cell analysis. *Mtb*-memory was assessed by subtracting the frequency of activated cell subsets inn unstimulated PBMC from *Mtb*-stimulated PBMC. **A**. Forest plot displays ln-transformed odds ratios (lnOR) and 95% confidence intervals (CI) of the differences mentioned above. The dotted line shows no average effect (lnOR = 0, corresponding to OR = 1). Features on the right side of this line are more highly expressed among LTBI-participants (OR > 1), and features on the left side are more highly expressed among TB-resisters (OR < 1). The table shows the absolute OR and the means of each parameter in TB-resisters and LTBI-participants. **B.** Typical scatter plots exemplifying differences in *Mtb*-memory CD8+IFNγ+ iNKT and between TB-resisters and LTBI-participants. **C**. Frequencies of CD8+IFNγ+ iNKT in all participants, including medians, upper and lower quartiles marked on the graph.

### Polyfunctionality of the immune cell subsets in TB-resisters and LTBI-participants

Polyfunctionality analyses based on Boolean gating of CD25, CD107a, GranzB, IFNγ, IL10 and PD1 in the T cell panel showed higher marker co-expression in TB-resisters than in LTBI-participants in *unstimulated* conditions on CD8+ iNKT, MR1-MAIT, CD8+ NKT and CD4+ and CD8+ Tconv (**Fig 4A**). In *Mtb-stimulated* conditions, CD8+ iNKT, MR1-MAIT, CD8+ NKT, and CD4+ and CD8+ Tconv maintained higher polyfunctionality in TB-resisters than LTBI-participants. LTBI-participants did not have any subsets with higher polyfunctionality than TB-resisters. There were no differences in polyfunctionality of NK and γδ T cells in Mtb-stimulated or unstimulated conditions between TB-resisters and LTBI-participants (**Fig S2**).

**Figure 4.**
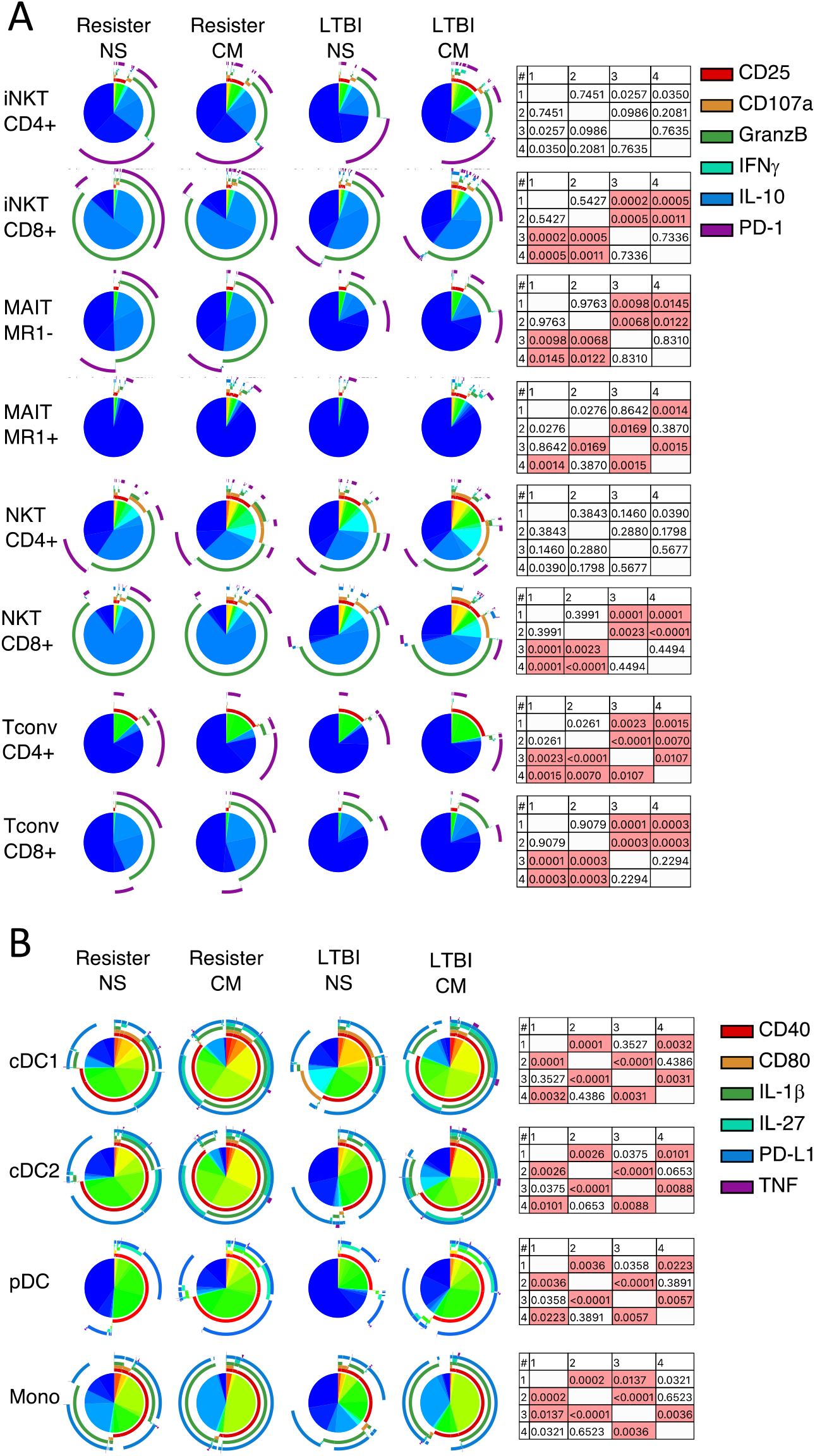
Polyfunctionality of immune responses in TB-resisters and LTBI-participants. Data were derived from 13 TB-resisters and 11 LTBI-participants. Pie charts 1-4 show proportions of cells expressing no markers (dark blue) or ≥1 marker (other colors) in response to *Mtb* stimulation or unstimulated control in the T cell panel (**A**) and APC panel (**B**). Arches show the markers expressed by the responding cells. The permutation tables present p-values for comparisons across TB-resisters unstimulated (1), TB-resisters *Mtb*-stimulated (2), LTBI-participants unstimulated (3) and LTBI-participants *Mtb*-stimulated (4) conditions. Tables show unadjusted p-values for each comparison. Pink boxes indicate significant differences defined by FDR p < 0.05.

In the APC analysis, we evaluated the expression of CD40, CD80, IL1β, IL27, PDL1 and TNFα on Mono, cDC1, cDC2, cDC total, and pDC. **Figure 4B** shows that TB-resisters had higher proportions of polyfunctional Mono than LTBI-participants in *unstimulated* conditions. No other differences were observed between TB-resisters and LTBI-participants.

### Coordination of the immune responses in TB-resisters and LTBI-participants

We complemented the analysis of manually gated PBMC subsets with Phenograph unbiased analysis, which identified 8 T-cell and 1 APC clusters that significantly differed between TB-resisters and LTBI-participants in *unstimulated* PBMC; 3 T-cell and 1 APC clusters in *Mtb-stimulated* conditions; and 5 T-cell and 3 APC *Mtb-memory* clusters (**Fig S3**). Most clusters confirmed findings already observed in the manually gated analysis, with the following exceptions: (1) Two T-cell and one APC *Mtb-memory* clusters (**Fig S3**) with higher relative frequencies in TB-resisters than LTBI-participants, in contrast to the manual gating analysis that did not identify excess *Mtb-memory* subsets in TB-resisters compared with LTBI-participants; and (2) A CD8+GMM+GranzB+ Tconv subset that had not been included in the prespecified manual gating analysis (**Fig 5 and S3**). This subset had higher frequency in *unstimulated* PBMC from TB-resisters compared with LTBI-participants (FDR-corrected p=0.008).

**Figure 5.**
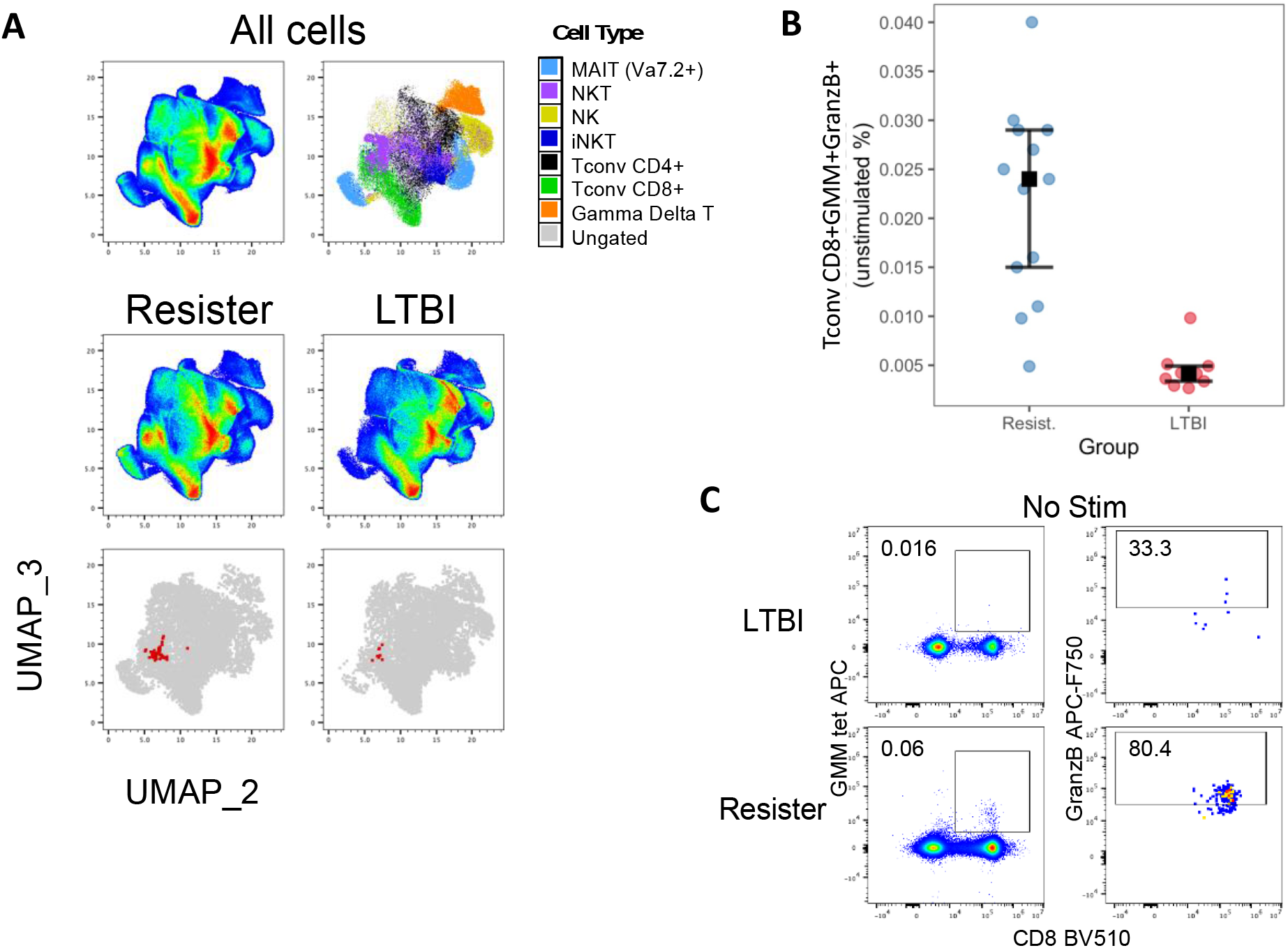
Comparison of CD8+GMM+GranzB+ Tconv subset in TB-resisters and LTBI-participants. Data were derived from 13 TB-resisters and 9 LTBI-participants that contributed 2,948,000 events down sampled to 67,000 events in the cleanup gate. **A**: T-cell UMAPs showing concatenated stimulated and unstimulated cells (upper left corner), distribution of major cell subsets (upper right corner); Distribution of unstimulated T cells in Resisters and LTBI-participants (middle row); CD8+GMM+GranzB+ cluster in Resister and LTBI-participants (bottom row). Representation of typical scatter plot. **B**: Individual results from each participant with medians, upper and lower quartiles are identified on the graph. **C:** Typical representation of the frequency of CD8+GMM+GranzB+ T cells in LTBI-participants and Resisters.

### Immune responses in BCG-recipients

The analysis of functional phenotypes in BCG-recipients showed significant differences with TB-resisters in 54 *unstimulated,* 41 *Mtb-stimulated,* and 41 *Mtb-memory* PBMC subset responses (**Fig S4**). Compared to LTBI-participants, BCG-recipients showed differences in 56 *unstimulated,* 32 *Mtb-stimulated,* and 24 *Mtb-memory* responses (**Fig S5**).

The comparison of polyfunctional NK and T cells in BCG-recipients and TB-resisters showed significantly higher frequencies of polyfunctional MR1+ and MR1-MAIT and CD4+ and CD8+ NKT and Tconv in TB-resisters both in *unstimulated* and *Mtb-stimulated* PBMC, while BCG-recipients had higher frequencies of polyfunctional CD4+ iNKT cells (**Fig 6A**). Compared with BCG-recipients, LTBI-participants had higher frequencies of polyfunctional MR1+ MAIT cells in *unstimulated* PBMC, while BCG-recipients had higher frequencies of CD4+ iNKT (**Fig 6B**). In *Mtb-stimulated* conditions, LTBI-participants showed higher frequencies of polyfunctional MR1+ MAIT and CD4+ Tconv, while BCG-recipients had higher frequencies of polyfunctional CD4+ and CD8+ iNKT.

**Figure 6.**
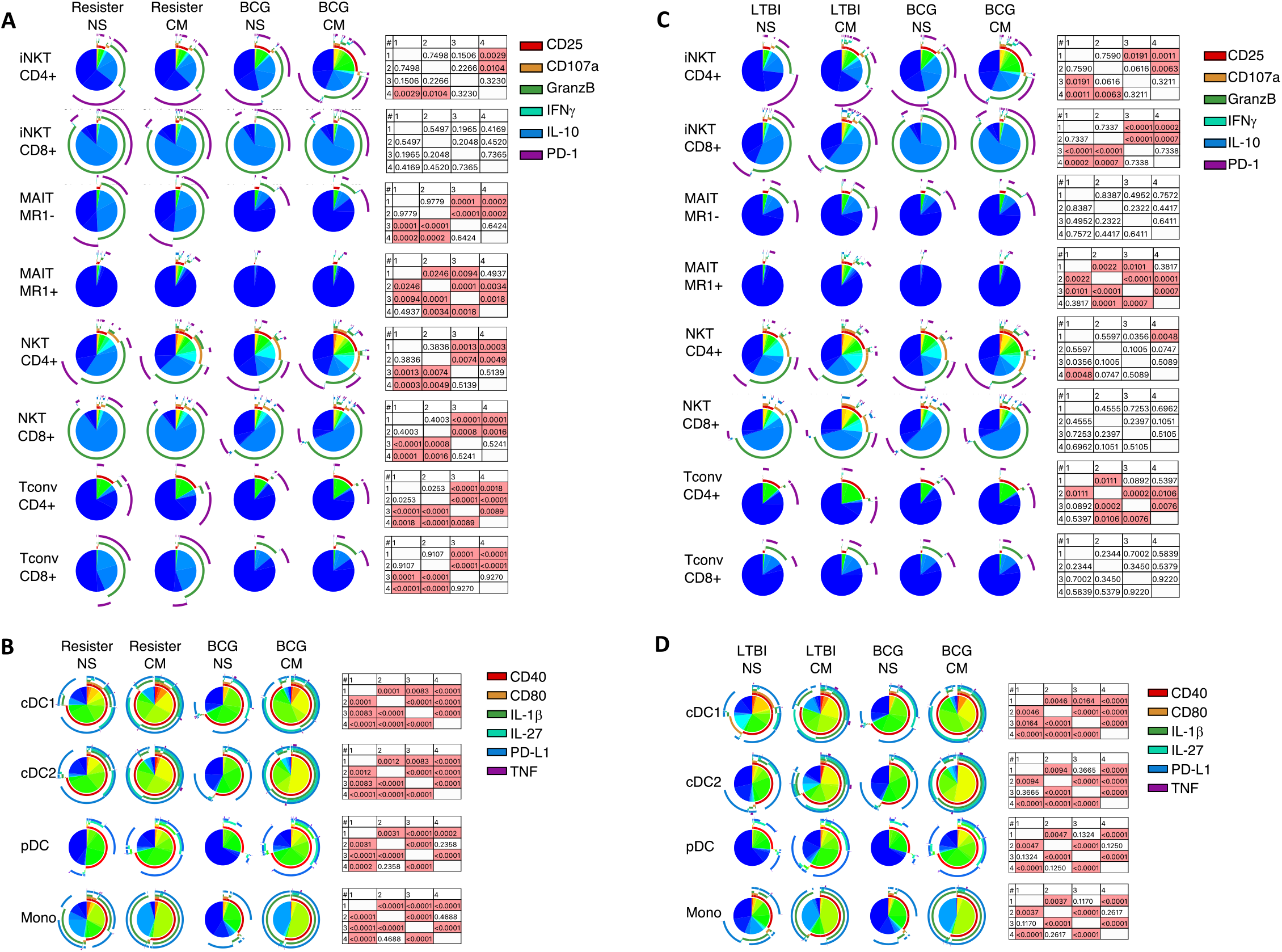
Polyfunctionality of immune responses in BCG-recipients by comparison with TB-resisters and LTBI-participants. Data were derived from 14 BCG-recipients, 13 TB-resisters, and 11 LTBI-participants. Pie charts 1-4 show proportions of cells expressing no markers (dark blue) or ≥1 marker (other colors) in response to *Mtb* stimulation or unstimulated control. Arches show the markers expressed by the responding cells. The permutation tables show unadjusted p-values for each comparison. Pink boxes indicate significant differences defined by FDR p < 0.05). **A:** T cell panel in BCG-recipients and TB-resisters; **B**: APC panel in BCG-recipients and TB-resisters; C: T cell panel in BCG-recipients and LTBI-participants; **D**: APC panel in BCG-recipients and LTBI-participants. The T cell panels show selected subsets with significant differences of BCG-recipients with TB-resisters and/or LTBI-participants defined by FDR p<0.05. Remaining subsets are shown in **Fig S6**.

The polyfunctionality analysis of APC showed that compared with BCG-recipients, TB-resisters had higher proportions of polyfunctional Monos, cDC1, cDC2 and pDC in *unstimulated* PBMC (**Fig 6C**). After *Mtb-stimulation* the cDC1 and cDC2 also had higher polyfunctionality in TB-resisters and none of the APC subsets had higher polyfunctionality in BCG-recipients. Compared to BCG-recipients LTBI-participants had higher frequencies of polyfunctional cDC1 in *unstimulated* PBMC, while BCG-recipients had higher frequencies of polyfunctional cDC1 and cDC2 after *Mtb stimulation* (**Fig 6D**).

One of the two recent descriptions of GMM+ Tconv showed their presence in response to BCG administration in infants, but not in adults^55^. On comparing the frequency of CD8+GMM+GranzB+ T cells in BCG-recipients with TB-resisters and LTBI-participants in our study, we also found very small frequencies of this cell subset (≤0.005% out of total lymphocytes) in BCG-recipients, significantly lower compared with either of the *Mtb*-exposed groups (FDR-corrected p≤0.00013; **Fig S7**).

## Discussion

The goal of this study was to characterize responses unique to TB-resisters compared to LTBI-participants that may be targeted by new TB vaccines. A consistent finding was increased GranzB-expressing conventional and nonconventional T cells in *Mtb-stimulated* and *unstimulated* PBMC of TB-resisters. This hallmark was observed in the prespecified manual gating and in the unsupervised cluster analyses and included expression of GranzB by Tconv, NKT and iNKT cells. The *Mtb-memory* analysis reveal higher cytotoxic responses in TB-resisters using the unbiased cluster analysis but not the manual gating. Collectively, these findings suggest that TB-resisters may mount higher and/or faster cytotoxic responses upon exposure to *Mtb* than LTBI-participants, which may contribute to the clearance of the infectious agent before establishing latency.

Of special interest was a population of CD8+GMM+GranzB+ Tconv that was not part of the prespecified manual gating but was identified by the Phenograph cluster analysis. This cell subset, which has only recently been described, was found in people with active TB, in whom CD8+GMM+ T cells displayed upregulated cytotoxic transcription profiles^56^. The authors concluded that this cell subset may contribute to the clearance of *Mtb*-infected host cells. Notably, we found higher proportions of CD8+GMM+GranzB+ T cells in TB-resisters compared with LTBI-participants, suggesting that this cell subset may contribute to the sterilizing immune protection of TB-resisters against *Mtb* infection.

TB-resisters also had higher proportions of PD1-expressing Tconv, iNKT and MR1-MAIT than LTBI-participants in *unstimulated* PBMC and after ex vivo *Mtb stimulation*. PD1 is an immunologic checkpoint inhibitor commonly expressed on activated T cells^57^. PD1 expression contributes to quenching the immune response after removal of the stimulating agent, limiting the inflammatory response and tissue destruction. Recent reports showed an association of increased T cell activation with TB morbidity and with increased risk of active *Mtb* infection in BCG-recipients^58–60^. Moreover, PD1 deficient mice as well as people treated with PD1/PDL1 blocking agents have increased risk of *Mtb* infection, morbidity, and reactivation^61–63^. Collectively, these observations suggest that the increased expression of PD1 in TB-resisters may limit their susceptibility to *Mtb* tissue destruction and, perhaps, propagation of infection.

A distinguishing characteristic of the immune response in TB-resisters from LTBI-participants was polyfunctionality. TB-resisters had higher proportions of polyfunctional Tconv and nonconventional T cells both in *Mtb-stimulated* and *unstimulated* conditions. Polyfunctional CD4+ Tconv, characterized by expression of IL2, IFNγ and/or TNFα have been previously associated with protection against active TB infection in animal models^64^. However, the role of polyfunctional CD4+ T cell responses in protection against *Mtb* infection in humans remains uncertain, particularly since a recent study showed that IL2, IFNγ and/or TNFα polyfunctional CD4+ T cell responses generated by BCG did not correlate with protection against TB disease in vaccinated infants^11^. In our study, cytokine production made a minor contribution to the polyfunctional T cell responses, which predominantly expressed GranzB, CD107a, CD25 and PD1, and involved mostly CD8+ T cells, including NKT, iNKT and Tconv. In addition, CD4+ Tconv and MR1-MAIT cells also showed higher polyfunctionality in TB-resisters than LTBI-participants expressing the same markers as the CD8+ Tconv. These findings suggest that high proportions of cytotoxic Tconv and nonconventional T cells may contribute to the immune protection of TB-resisters against *Mtb* infection.

*Mtb-memory* T cell responses identified by the expression of single activation markers were higher in LTBI-participants than in TB-resisters, although two *Mtb-memory* CD4+ Tconv clusters revealed by the Phenograph analysis had higher frequencies in TB-resisters than in LTBI-participants. These data confirm that TB-resisters develop *Mtb*-specific T cell memory, but in general *Mtb-memory* responses are higher in LTBI-participants. This finding may be related to the observation that 25% of people with LTBI have evidence of active *Mtb* replication, which continuously exposes the immune system to *Mtb* antigens and maintains high levels of memory cells^65–68^.

APC displayed few differentiating features between TB-resisters and LTBI-participants. In *unstimulated* PBMC, TB-resisters had higher frequencies of activated and polyfunctional Mono by manual gating and Boolean analyses, respectively, and cDC2 by cluster analysis. However, no differences were found in *Mtb-stimulated* responses between TB-resisters and LTBI-participants. It is important to note that *Mtb* contains multiple toll-like receptor (TLR) agonists, which hinders the discrimination of *Mtb-stimulated* responses from nonspecific stimulation TLR-mediated activation of APC. Nevertheless, the Phenograph analysis identified an activated Mono cluster with higher frequency in TB-resisters than LTBI-participants.

The analysis of *Mtb-memory* responses in BCG-recipients using single markers of activation showed both higher and lower responses in multiple T-cell and APC subsets compared with TB-resisters or LTBI-participants. These differences may have been related to the difference in environmental exposures, including *Mtb*, in addition to genetic backgrounds of BCG-recipients compared with TB-resisters or LTBI-participants. The Boolean analysis of polyfunctional responses narrowed down the differences between BCG-recipients and LTBI-participants or TB-resisters. Overall, BCG-recipients had higher frequencies of *unstimulated* and *Mtb-stimulated* iNKT polyfunctional cells than both LTBI-participants and TB-resisters. BCG-recipients also had lower frequencies of *unstimulated* and/or *Mtb-stimulated* Tconv, γδ, NKT, and/or MAIT polyfunctional cells compared to both LTBI-participants and TB-resisters, but a larger number of polyfunctional subsets differentiated BCG-recipients from TB-resisters than LTBI-participants. Compared to TB-resisters, BCG-recipients also had universally lower frequencies of *stimulated* and *Mtb-stimulated* polyfunctional APC subsets. Compared to LTBI-participants, BCG-recipients had lower frequencies only of *stimulated* and *Mtb-stimulated* polyfunctional cDC1 and higher frequencies of polyfunctional cDC2. Collectively, these findings suggest superior polyfunctionality of T cells and APC in TB-resisters compared with BCG-recipients.

Our study has both strengths and limitations. The main limitations consisted of the small number of participants and different demographic features of BCG-recipients compared with C-TRIUMPH participants (e.g., race/ethnicity). Although our study did not measure antibody responses, which have been recently shown to differentiate TB-resisters from LTBI-participants^17,69^, the CMI analysis was comprehensive and revealed new findings unique in TB-resisters.

We conclude that both innate and adaptive responses distinguish TB-resisters from LTBI-participants. Unique elements of the cell-mediated immune responses to *Mtb* in TB-resisters are high cytotoxic potential, expression of immunologic checkpoint inhibitors and polyfunctionality of Tconv and nonconventional T cells, suggesting that some or all may be key factors in an effective immune response against *Mtb*. Although TB-resisters displayed these characteristics even without ex vivo *Mtb stimulation*, it is reasonable to propose that a vaccine that increases some or all these responses in the form of *Mtb-memory* may be more effective than BCG.

## Materials and Methods

### Participants

Cryopreserved PBMC were obtained from 13 TB-resisters and 11 LTBI-participants in C-TRIUMPH ^23,52,53^. LTBI-participants were defined by positive IGRA and/or TST at entry. TB-resisters had negative IGRA and TST at entry and over the 2 years of follow up. BCG-recipients whose immunization status was documented by medical history, who lived most of their lives outside of TB endemic areas, and who had negative IGRA were enrolled at the University of Colorado Anschutz Medical Campus (CU-AMC). <u>IGRA</u> testing used Quantiferon Gold in Tube kit (Qiagen) performed as per manufacturer’s instructions.

### Flow Cytometry

Testing conditions were optimized using a convenience sample of PBMC from LTBI-participants and donors without previous exposure to TB or BCG submitted to stimulation for several time intervals with different concentrations of irradiated *Mtb,* whole cell lysate, cell membrane, PPD and BCG. *Mtb* cell membrane provided the best discrimination between LTBI-participants and TB-unexposed and was selected for this study. PBMC were thawed in serum-free media (AIM V, Gibco) supplemented with benzonase (50 units/mL, Millipore). PBMC were resuspended at 2*10^6^ cells/ml and stimulated with cell membrane (BEI Resources Strain CDC1551, Cell Membrane Fraction, NR-14832) at 20ug/ml in AIM V or unstimulated overnight. Brefeldin-A and Monensin (SIGMA) at 5ug/ml each were added for the last 4h of culture. For the T-cell panel, CD107a-BV785 (Biolegend) was added for the last 4h of culture. Cells were washed with PBS, stained with Zombie NIR Fixable Viability dye (Biolegend), and washed with PBS+1% albumin. For the APC panel, cells were treated with Human TruStain FcX (Biolegend), surface-stained with CD16-eFluor450, CD83-PerCP-eFluor710 (Thermo Fisher/eBioscience), CD40-BV480, CD141-BV750, CD11c-BB515 (BD Biosciences), CD80-BV510, CD1c-BV605, CD123-BV650, PDL1-BV785, HLA-DR-AlexaFluor488, CD3-, CD19-, CD20- and CD56-PerCP-Cy5.5, CD14-APC-Fire750 (Biolegend), brilliant stain buffer plus (BD Biosciences) and True-stain monocyte blocker (Biolegend). Cells were washed and treated with eBioscience Foxp3 Staining Buffer Set (Thermo Fisher/eBioscience), then stained with IL8-BV421, IL10-BV711, IL1β-PE (BD Biosciences), GMCSF-PE-Dazzle594, IL12p40-AlexaFluor647 (Biolegend), IL27-APC, TNFα-AlexaFluor700 (Thermo Fisher/eBioscience) and brilliant stain buffer plus. For the T-cell panel, cells were treated with 50% human AB serum (Gemini) in PBS+1% albumin, washed and surface-stained with tetramers: CD1b-unloaded-FITC, CD1b-GMM-APC, and MR1-BV421. Cells were washed, stained with TCR Vα7.2-BV605, TCR Vα24-Jα18-PE-Cy7 (Biolegend) and TCR γδ-PE (BD Biosciences) followed by CD16-eFluor450, CD3-AlexaFluor532 (Thermo Fisher/eBioscience), CD56-BV480, CD25-BB515 (BD Biosciences), CD8-BV510, PD1-BV711, CD69-PE-Cy5, CD161-AlexaFluor700 (Biolegend) and brilliant stain buffer plus. Cells were washed, treated with Foxp3 Staining Buffer Set, then stained with IL17-BV570, IL2-BV650, IL10-PE-Dazzle594, GranzymeB-APC-Fire750 (Biolegend), IFNγ-BV750, GMCSF-AlexaFluor647 (BD Biosciences), Ki67-PerCP-eFluor710, TNFα-PerCP-Cy5.5 (Thermo Fisher/eBioscience) and brilliant stain buffer plus. Cells from both panels were washed and fixed with PBS+1% paraformaldehyde, acquired on the Cytek Aurora™ (Cytek Biosciences), and analyzed with FlowJo (Becton Dickinson). Two leucopack controls were used in each run to ensure inter-run reproducibility and to examine technical effects.

### Boolean analysis

Boolean combinatorial gates were created using the 6 cytokines/activation markers, generating 64 distinct activation phenotypes. Graphical representation was performed using SPICE 6.0 software^70^. The data were analyzed using permutations (10,000 iterations) included in the software.

### UMAP analysis

For T cell clustering, we used 67,000 down-sampled events from 13 TB-resisters and 9 LTBI-participants for a total of 2,948,000 events in the concatenated file. For APC clustering, we used data from 11 TB-resisters and 8 LTBI-participants for a total of 502,479 events in the concatenated file. Parameters were rescaled to ArcSinh prior to running UMAP with phenograph in FlowJo on the concatenated file using parameters, Euclidean distance, 15 nearest neighbors and minimum distance of 0.5^71^.

### Statistical Testing of Cell Lineage and Function Between TB-Resisters and LTBI-participants

To compare the average cell lineage frequencies obtained from manual and data-driven gating between groups, we used beta regression *(betareg* R package, v3.1-3) with a logit-link to model cell lineage percent (regression outcome; between 0 and 100% of events) on group (primary explanatory variable of interest; LTBI-participant versus TB-resisters), adjusting for participant age in years as a covariate ^72^. Similarly, to test for functional differences among *unstimulated* conditions, we modeled percent positivity (outcome; between 0 and 100% of cells in given lineage expressing functional marker) on group, correcting for age. “Rare” outcomes (i.e., outcomes of 0% in over 1/3^rd^ of participants) were excluded from testing due to their low variability among our participants. For each set of statistical tests, we defined significant between-group differences as a non-zero group regression coefficient (Wald test, multiple testing adjustment using the Benjamini-Hochberg false discovery rate < 0.05)^73^.

To examine differences in functional markers under *Mtb-stimulated* conditions, we repeated this modeling procedure but with the stimulated percent positivity values as the regression outcome variable, adjusting for age. We also examined differences in *Mtb-memory* by additionally adjusting for each participant’s corresponding baseline values (unstimulated percent positivity values in their paired control sample) as a covariate. The *Mtb-memory* analysis thus emphasizes markers that were not already differentially abundant between groups prior to stimulation, or stimulation effects potentially masked by unstimulated differences. Adjusting for baseline levels are conceptually similar to “subtracting” the unstimulated values from the Mtb-stimulated values; as a consequence, we visualize Mtb-memory with Δ-values, defined by stimulated frequencies minus unstimulated frequencies. Functional markers with limited changes following Mtb-stimulation (i.e., Δ-values of magnitude ≤ 0.1% in over 1/3^rd^ of participants) were excluded from testing due to their low variability among our participants and tendency to recapitulate unstimulated tests. Significant between-group differences in Mtb-memory were based on FDR < 0.05.

These modeling procedures was also repeated for the TB-resister versus BCG recipient comparisons and the LTBI-participants versus BCG recipient comparisons. All analyses were performed in the R statistical package v4.0.2 unless otherwise specified. Additional methodical details and analysis code is publicly available at https://github.com/chooliu/Sterile_vs_Latent_TB_Immunity.

## Supporting information

Manuscript

## Declaration of approval

The study was approved by the Colorado Multiple Institutional Review Board (COMIRB) and by the IRBs of the Byramjee Jeejeebhoy Govt. Medical College, India and Johns Hopkins Medicine. All participants signed informed consent.

## Author contributions

AW designed the study, oversaw the laboratory assays, analyzed the data and prepared the manuscript. EJ developed the flow cytometry assays, analyzed the data and contributed to manuscript preparation. CL performed the statistical analysis and contributed to manuscript preparation. NL conducted regulatory and clinical activities at CU-AMC. For the clinical research cohort activities in Pune, India: VM, MP, RL, CV oversaw the clinical enrollment and follow-up activities; VK, RB, AK conducted the laboratory activities and specimen management; NG conducted the data management activities; MP oversaw all the day-to-day study coordination activities; and AG, VM obtained the funding for the clinical cohort protocol, and AG, VM, and MP developed the cohort protocol and monitored the overall study conduct at BJGMC JHU CRS. All authors reviewed and approved the manuscript.

## Acknowledgments

The authors thank Ms. Sarah Lazarus for technical support; Drs. Seshadri and Moody for technological and material support on GEMT tetramers^74^ and GMM, respectively; and the NIH tetramer core for MR1 tetramers and for CD1b monomers used for generation of GEMT tetramers. The MR1 tetramer technology was developed jointly by Dr. James McCluskey, Jamie Rossjohn, and David Fairlie^75^, and the material was produced by the NIH Tetramer Core Facility as permitted to be distributed by the University of Melbourne. The following reagent was obtained through BEI Resources, NIAID, NIH: *Mycobacterium tuberculosis,* Strain CDC1551, Cell Membrane Fraction, NR-14832.

## Funding

The study was supported by NIAID grant 1R21AI134129 (AW). CTRIUMPH is part of the Regional Prospective Observational Research for Tuberculosis consortium for India funded with Federal funds from the Government of India’s (GOI) Department of Biotechnology (DBT), the Indian Council of Medical Research (ICMR), the USA National Institutes of Health (NIH), the National Institute of Allergy and Infectious Diseases (NIAID), the Office of AIDS Research (OAR), and distributed in part by CRDF Global (USB1-31147-XX-13 to AG). This work was also supported by the NIH funded Johns Hopkins Baltimore-Washington-India Clinical Trials Unit for NIAID Networks (UM1AI069465 to AG). Rahul Lokhande and Chhaya Valvi were supported by the BJGMC JHU HIV TB Program funded by the Fogarty International Center, NIH (D43TW009574). The contents of this publication are solely the responsibility of the authors and do not represent the official views of the DBT, the ICMR, the NIH, or CRDF Global. Any mention of trade names, commercial projects or organizations does not imply endorsement by any of the sponsoring organizations. The funders had no role in study design, data collection and analysis, decision to publish, or preparation of the manuscript.

## Data and materials availability

The flow cytometry was partially uploaded to ImmPort (https://www.immport.org) under study accession SDY1694. Information about C-TRIUMPH can be found at https://main.ccghe.net/content/report-c-triumph-cohort-tuberculosis-research-indo-us-medical-partnership.

## Notes

**Conflicts of interest**: AW receives research grants from GlaxoSmithKline and Merck (moneys to CU) and personal fees from Merck and Sequirus. No other conflicts exist.

### Competing Interest Statement

The authors have declared no competing interest.

### Summary of Updates

Correct figure labeling

