## Supplementary material for "Comparative immune responses to *Mycobacterium tuberculosis* in people with latent infection or sterilizing protection": Manuscript

**APCTable S1. Flow cytometry panels**

| **Panel** | **Cell type** | **Lineage markers** | **Functional markers** |
| --- | --- | --- | --- |
| **T cells** | Tconv | CD3^+^TCRγδ^-^ Vα24-Jα18^-^Vα7.2^-^ CD56^-^ | CD25, CD69, CD107a, GranzB, GMCSF, IL2, IL10, IL17, IFNγ, TNFα, Ki67, and PD1 |
|  | *NK* | CD3^-^CD16^+^ and/or CD56^+^ |  |
|  | *γδT* | CD3^+^TCRγδ^+^ |  |
|  | *iNKT* | CD3^+^Vα24-Jα18^+^ |  |
|  | *NKT* | CD3^+^TCRγδ^-^ Vα24-Jα18^-^Vα7.2^-^CD56^+^ |  |
|  | *MAIT* | CD3^+^TCR Vα7.2^+^MR1-tetramer^+/-^ |  |
|  | *GEMT* | CD3^+^Vα7.2^+^GMM-CD1b-tetramer^+^ |  |
| **APC** | *Monocyte* | Lin^-^HLADR^+^CD14^+^CD16^-/+^ | CD40, CD80, CD83, IL1β, IL8, IL10, IL12p40, IL27, TNFα, PDL1 |
|  | *cDC1* | Lin^-^HLADR^+^CD14^-^CD11c^+^CD141^+^CD1c^-^ |  |
|  | *cDC2* | Lin^-^HLADR^+^CD14^-^CD11c^+^CD141^-^CD1c^+^ |  |
|  | *pDC* | Lin^-^HLADR^+^CD14^-^CD11c^-^CD123^+^ |  |

Abbreviations: cDC= conventional dendritic cells; pDC= plasmacytoid DC; mono= monocytes; GEMT= germ-line encoded, mycoyl-reactive T cells; iNKT= invariant natural killer T cells; MAIT=mucosal-associated invariant T cells; NK= natural killer cells; NKT=natural killer T cells; Tconv=conventional T cells.

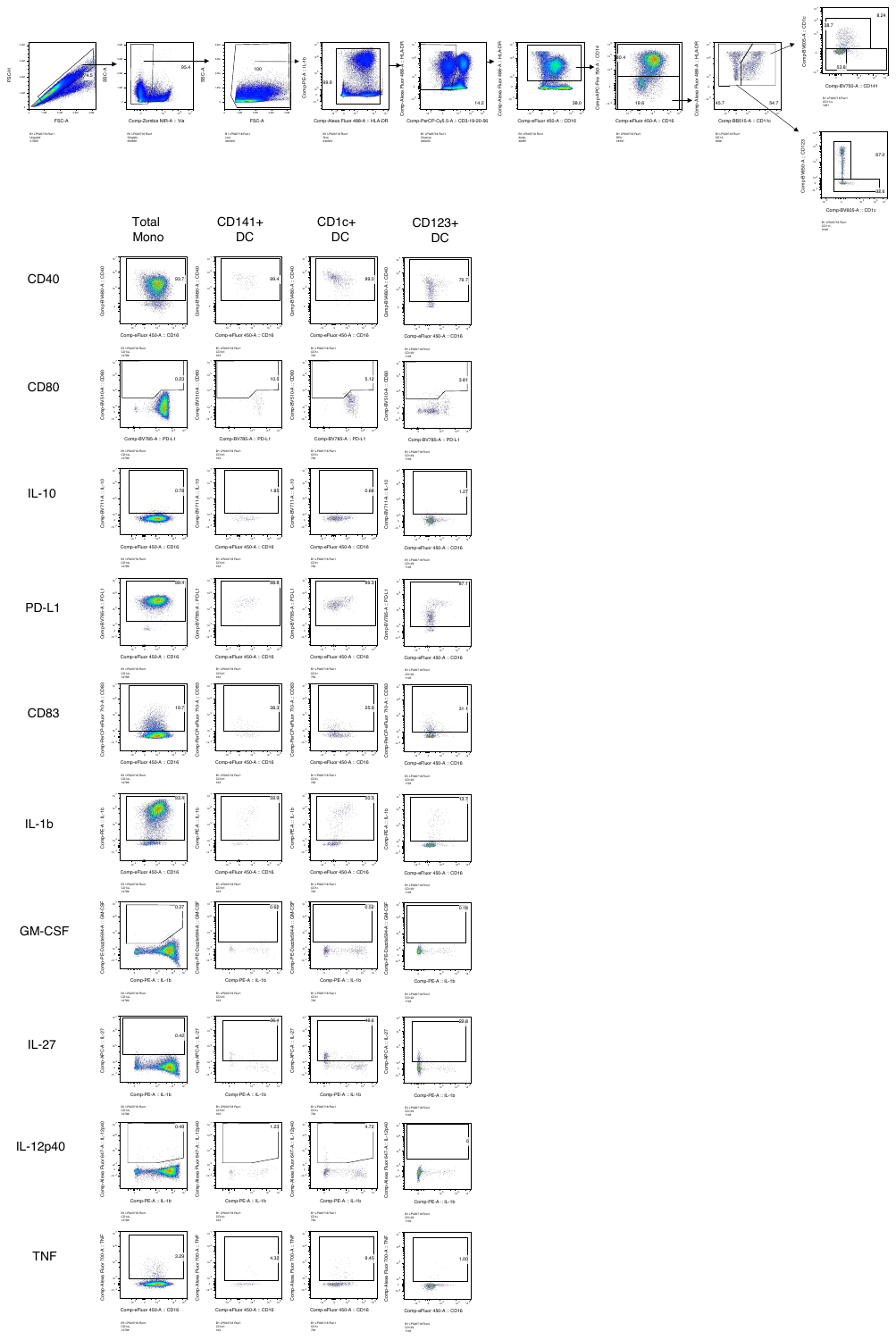

APC

**Figure S1. Gating strategy**. **Top**: T cells including NK cells. **Bottom**: APC.

GD

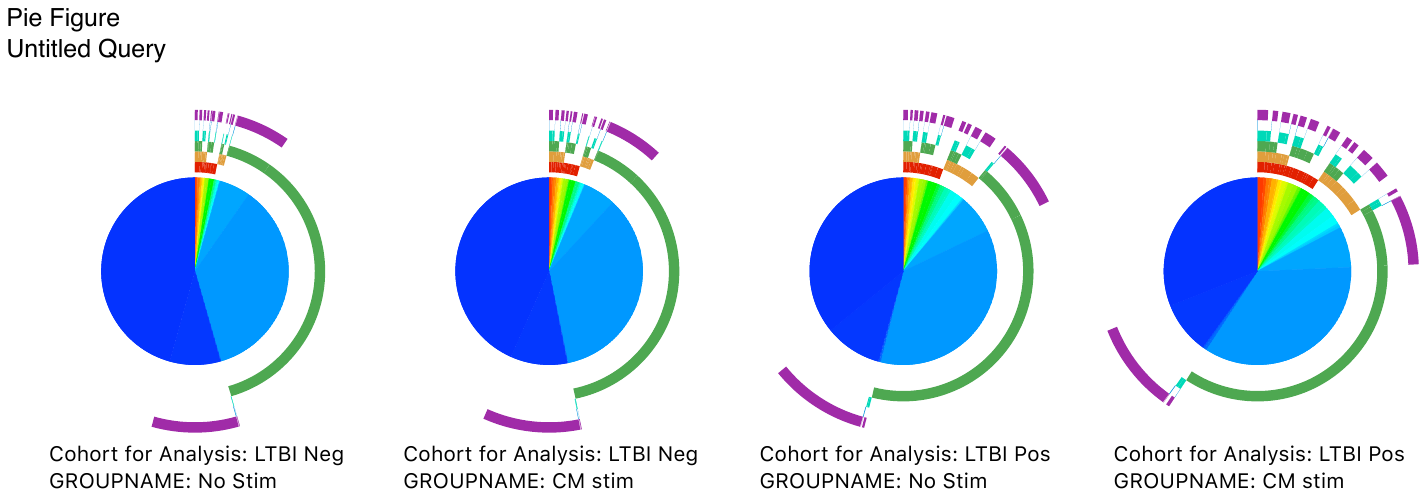

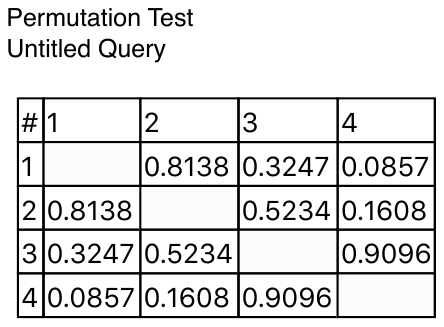

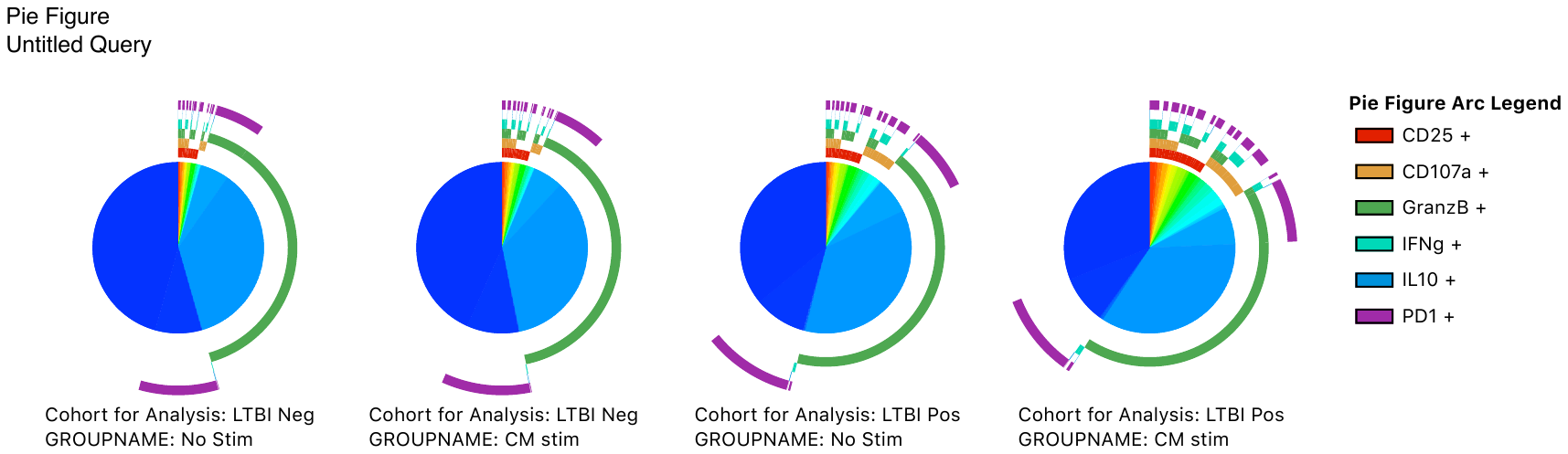

CD25

CD107a

GranzB

IFNγ

IL-10

PD-1

NK

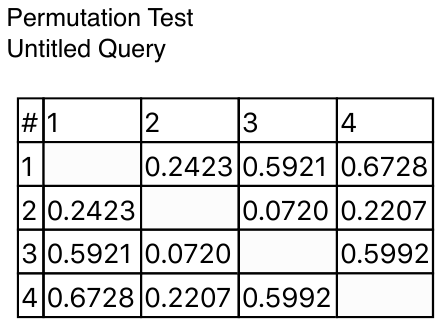

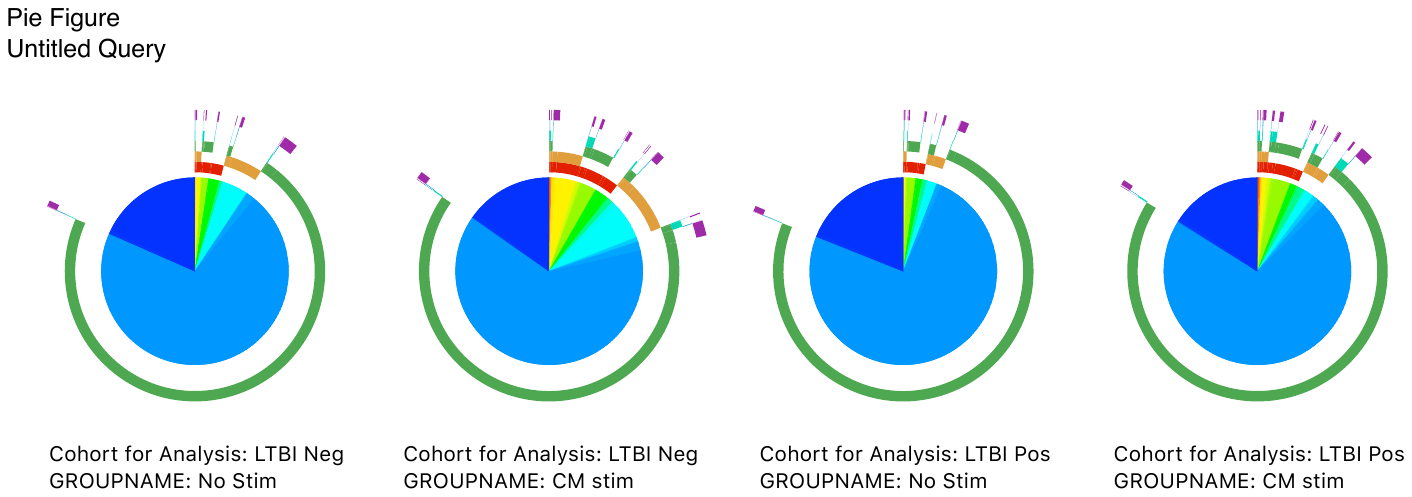

LTBIneg

NS

LTBIneg

CM

LTBIpos

NS

LTBIpos

CM

**Figure S2. Polyfunctionality of the immune responses to Mtb in TB-resisters (LTBIneg) compared with LTBI-participants (LTBIpos).** Data were derived from 13 TB-resisters and 11 LTBI+ participants. Pie charts 1-4 show proportions of cells expressing no markers (dark blue) or ≥1 marker (other colors) in response to Mtb stimulation or control. Arches show the markers expressed by the responding cells. The permutation tables present p values for comparisons across TB-resisters unstimulated (1), TB-resisters Mtb-stimulated (2), LTBI+ participants unstimulated (3) and LTBI+ participants Mtb-stimulated (4) conditions. Tables show unadjusted p-values for each comparison. Pink boxes indicate significant differences defined by FDR p < 0.05.

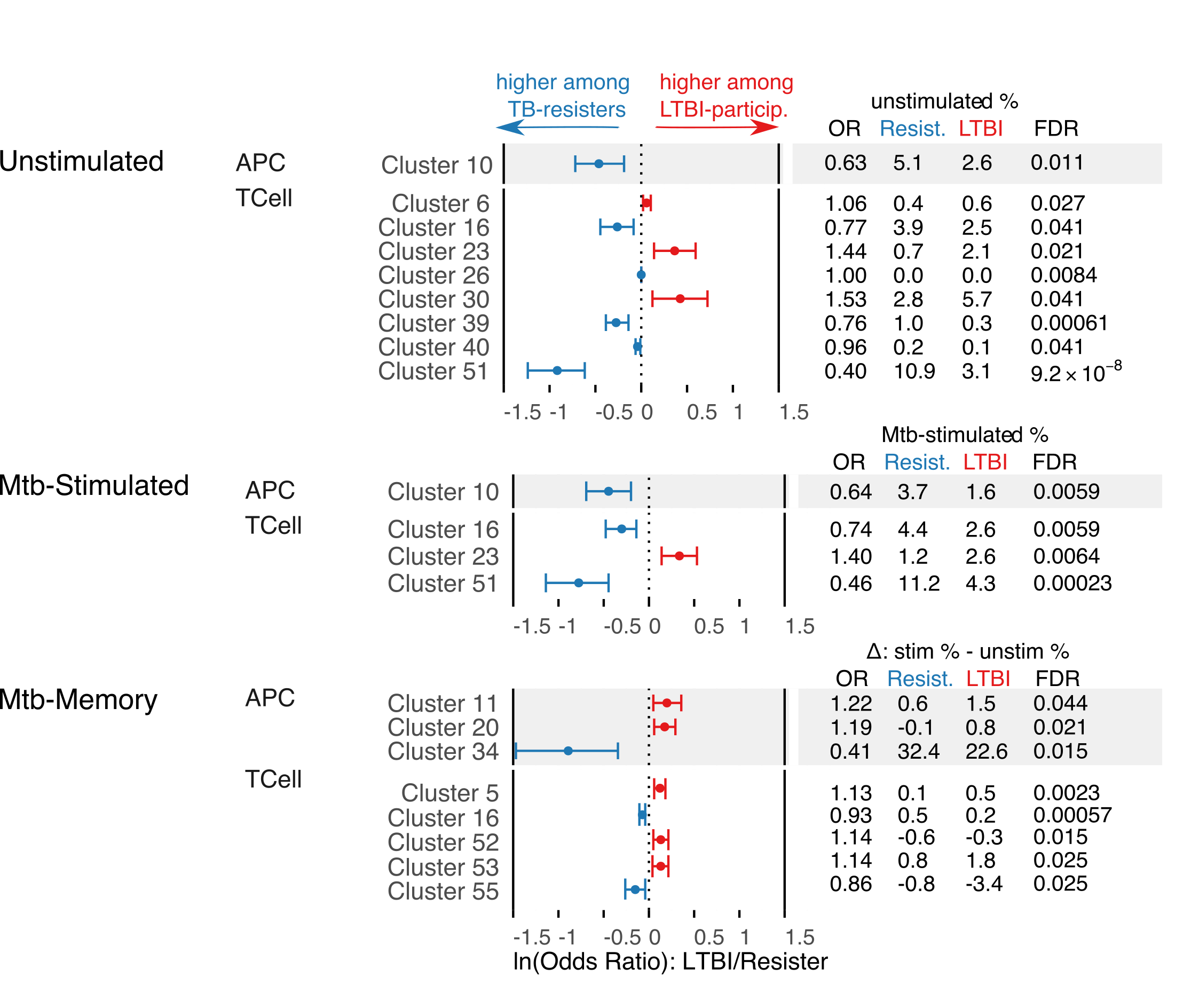

**A**

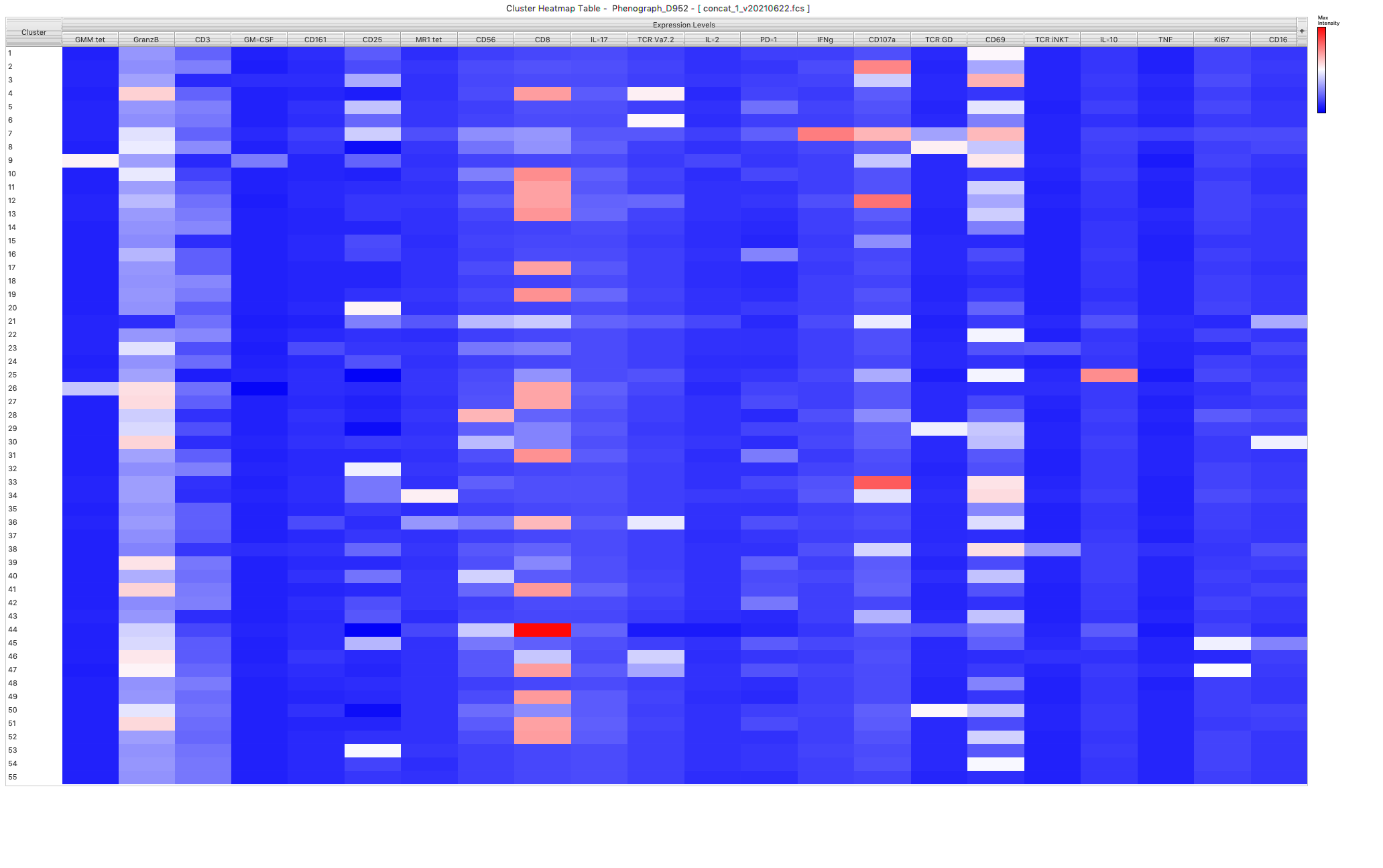

**B**

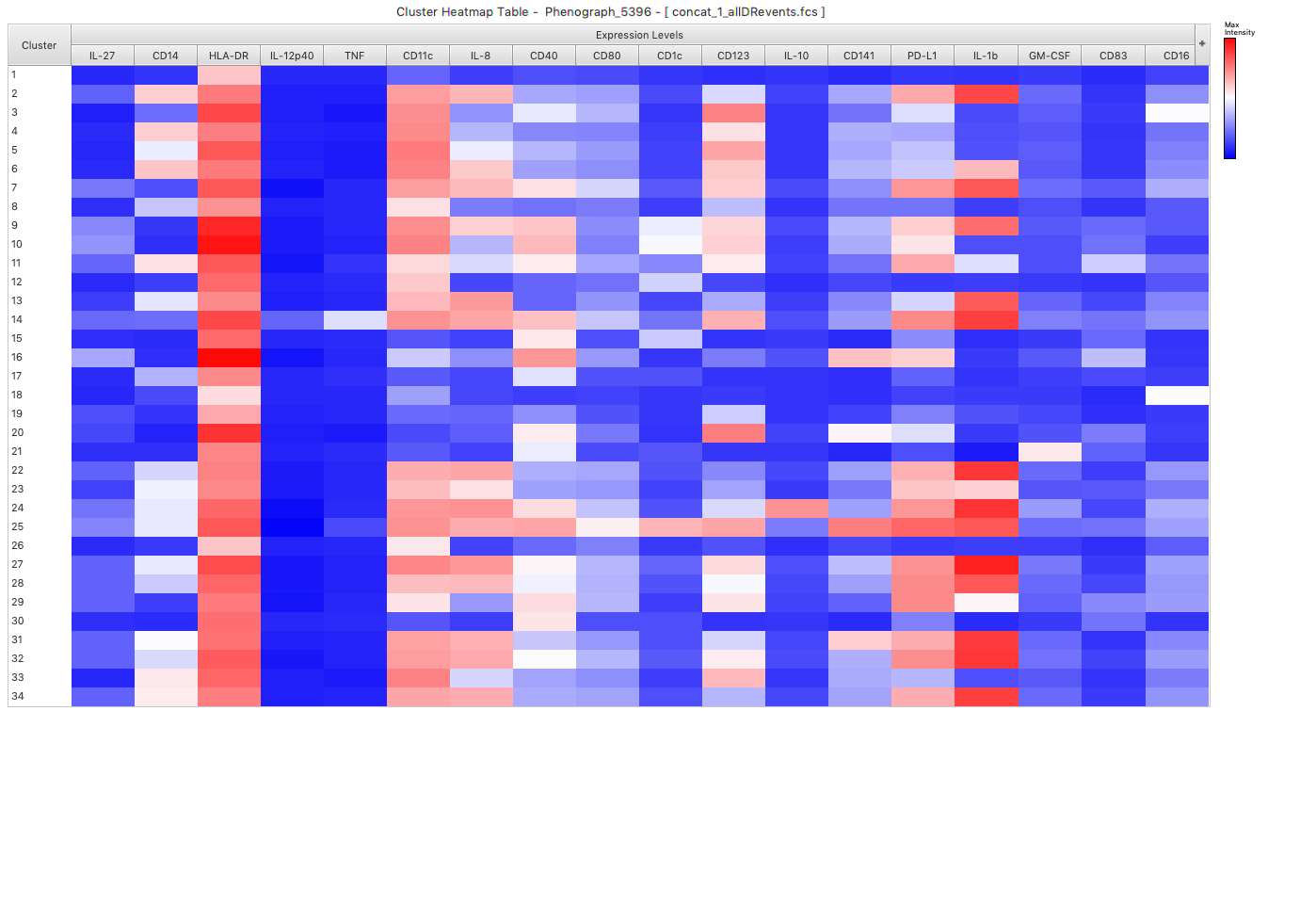

**C**

**Fig S3. Cluster analysis of adaptive and innate immune responses in TB-resisters and LTBI-participants.** Data were derived from 13 TB-resisters and 9 LTBI-participants for the T cell clustering and 12 TB-resisters and 8 LTBI-participants for the APC clustering. **A**: Forest trees of lnOR and 95% CI, means, and FDR-corrected p values of clusters significantly different between the two groups. **B**: Heat map of T-cell clusters identified by Phenograph. **C**: Heat map of APC clusters.

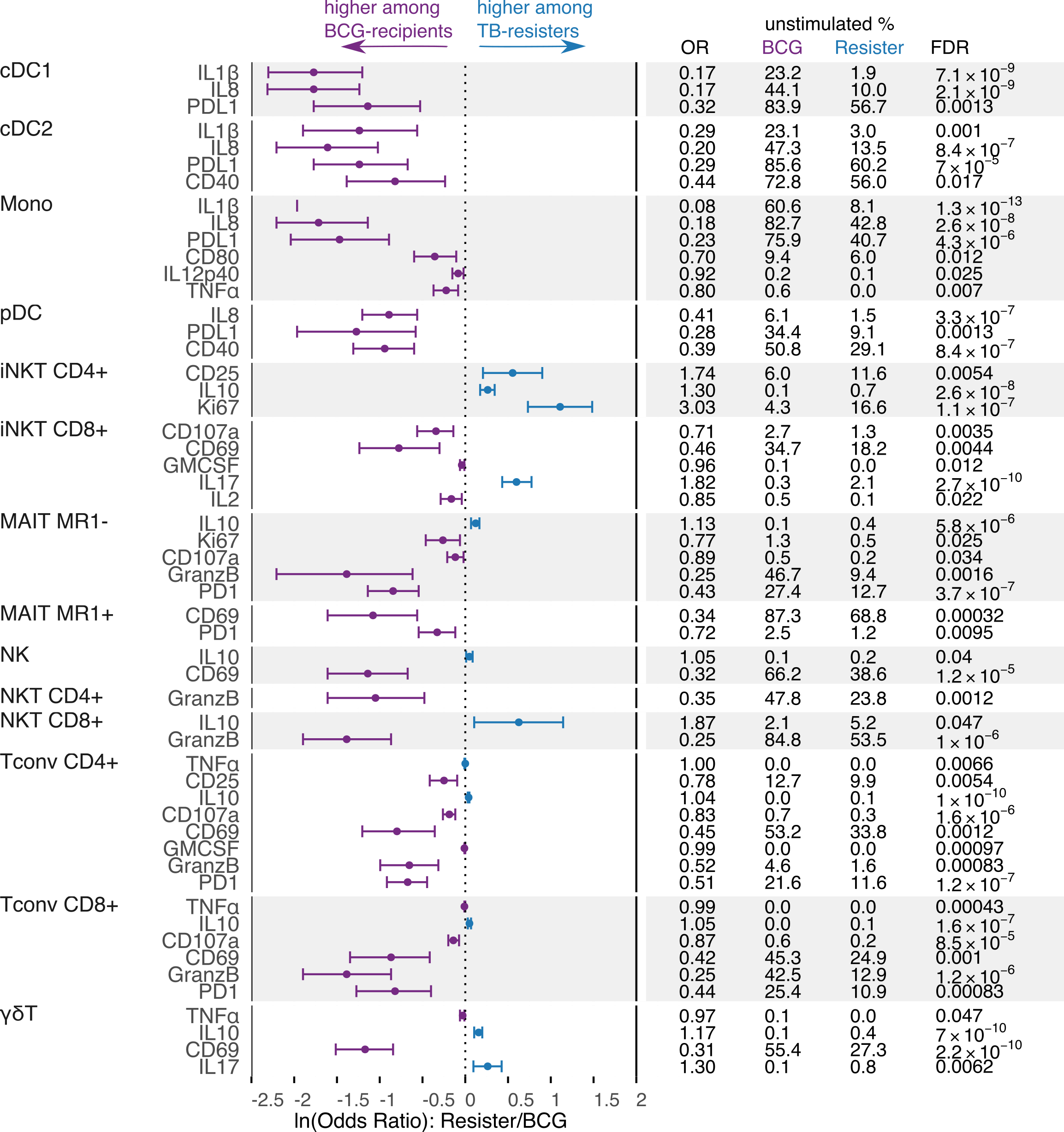

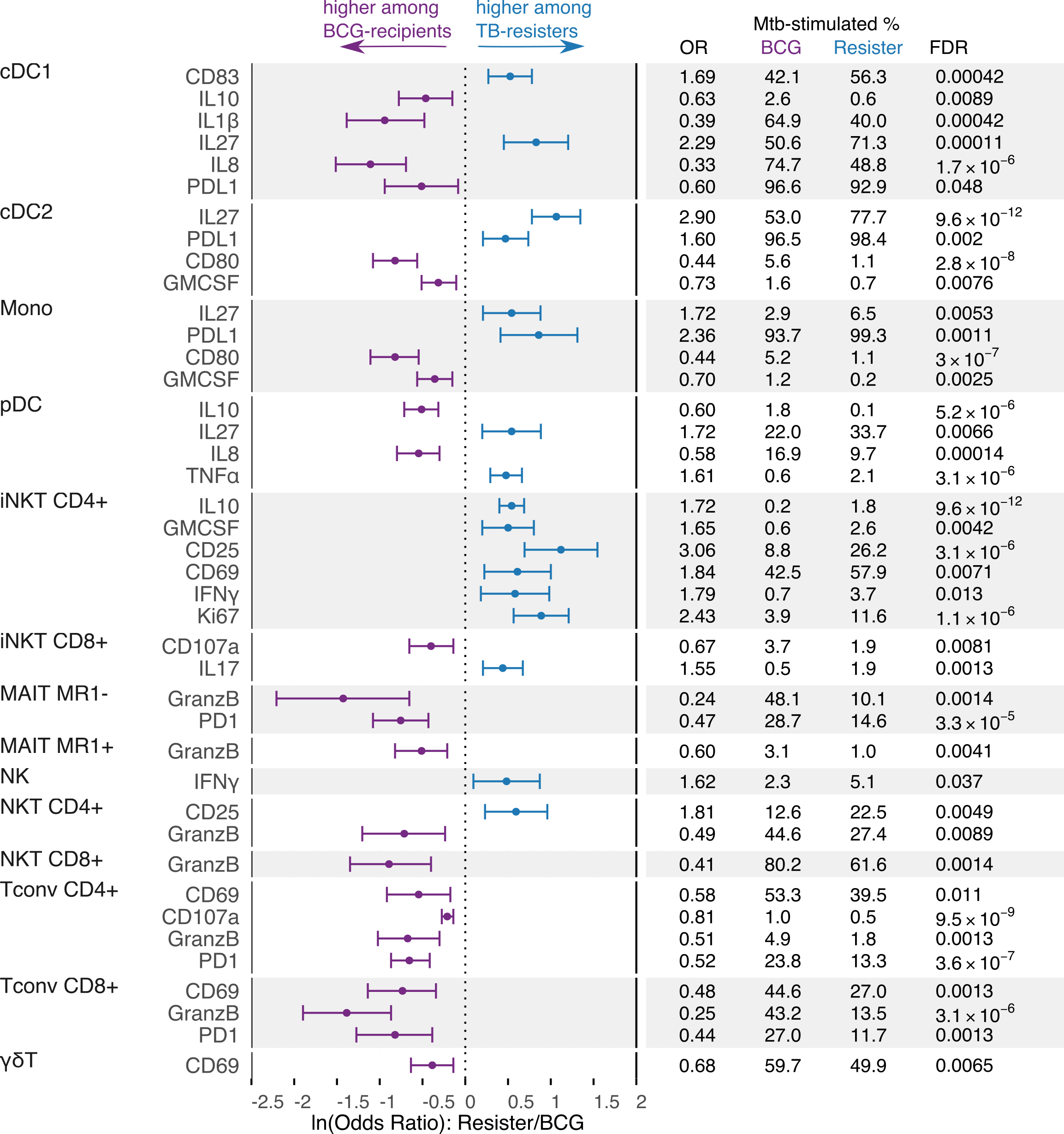

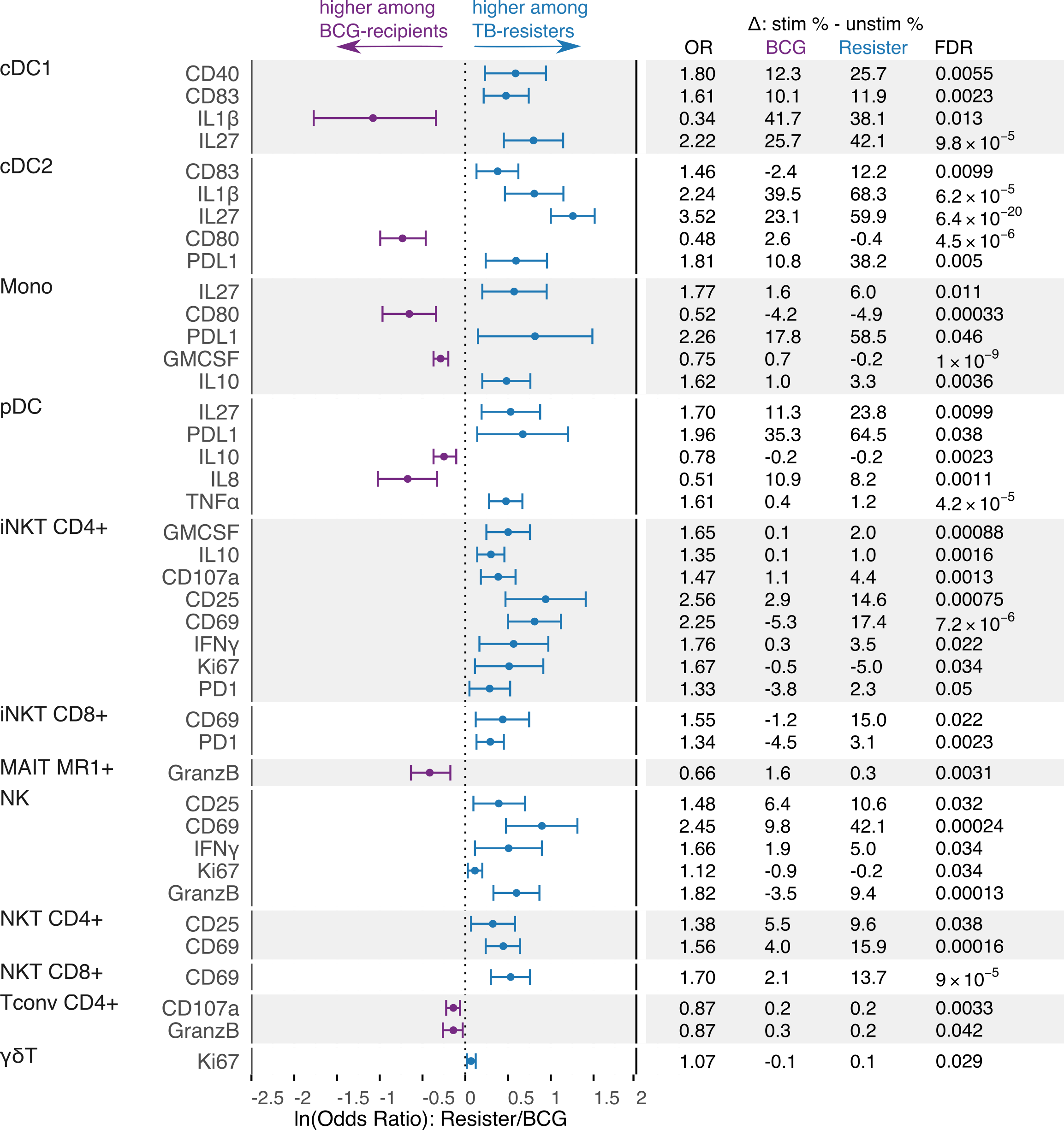

**Figure S4**. **Comparison of immune cell subset frequencies in BCG recipients and TB-Resisters.** Data were derived from 13 TB-Resisters and 14 BCG recipients. Forest plot displays ln-transformed odds ratios (lnOR) and 95% confidence intervals (CI). The dotted line shows no average effect (lnOR = 0, corresponding to OR = 1). Features on the right side of this line are more highly expressed among TB-Resisters (OR > 1), and features on the left side are more highly expressed among BCG recipients (OR < 1). The table shows the absolute OR and the means of each parameter in TB-resisters and LTBI-participants. **Top**: unstimulated PBMC; **Middle**: Mtb-stimulated; **Bottom**: Mtb-memory.

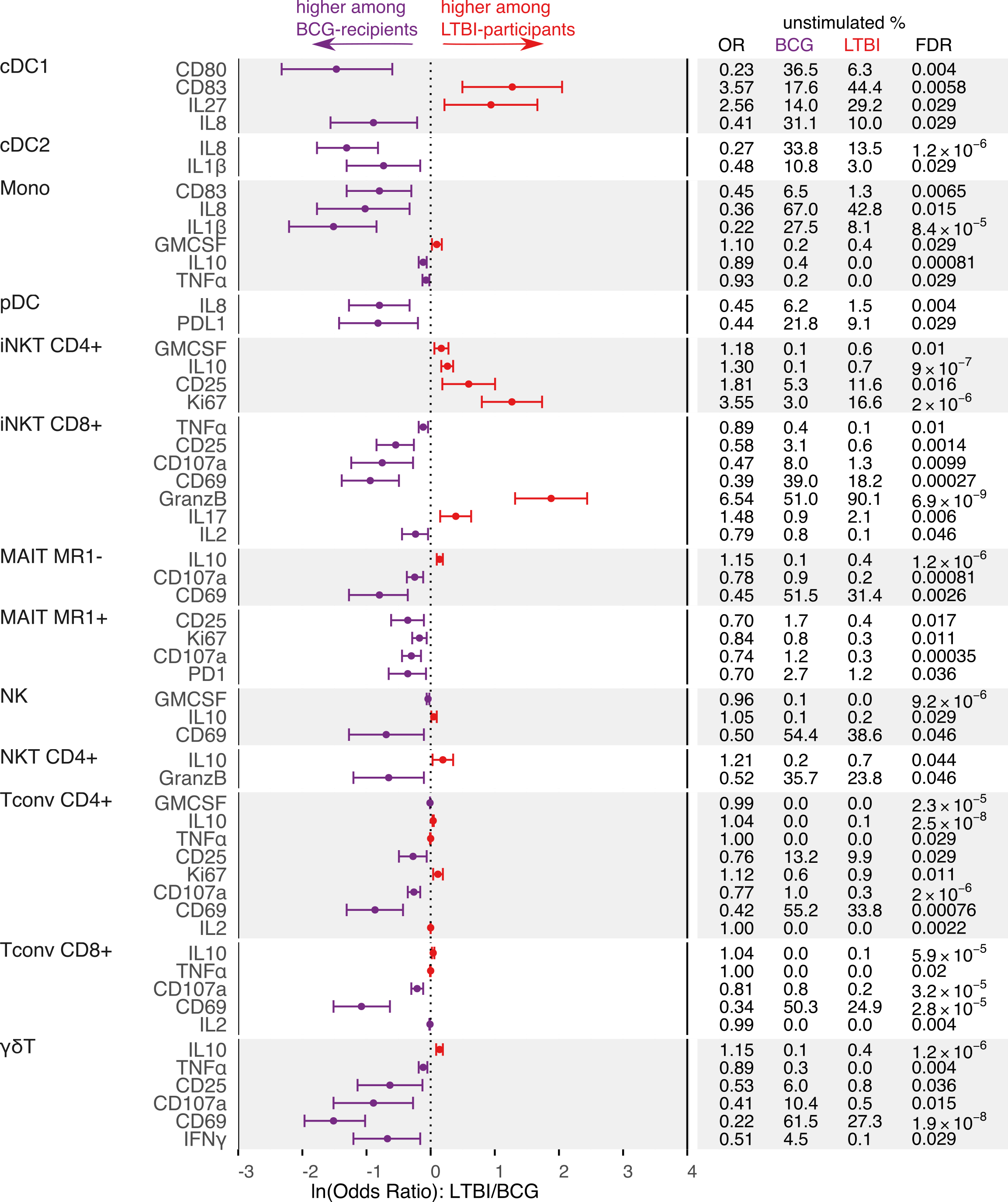

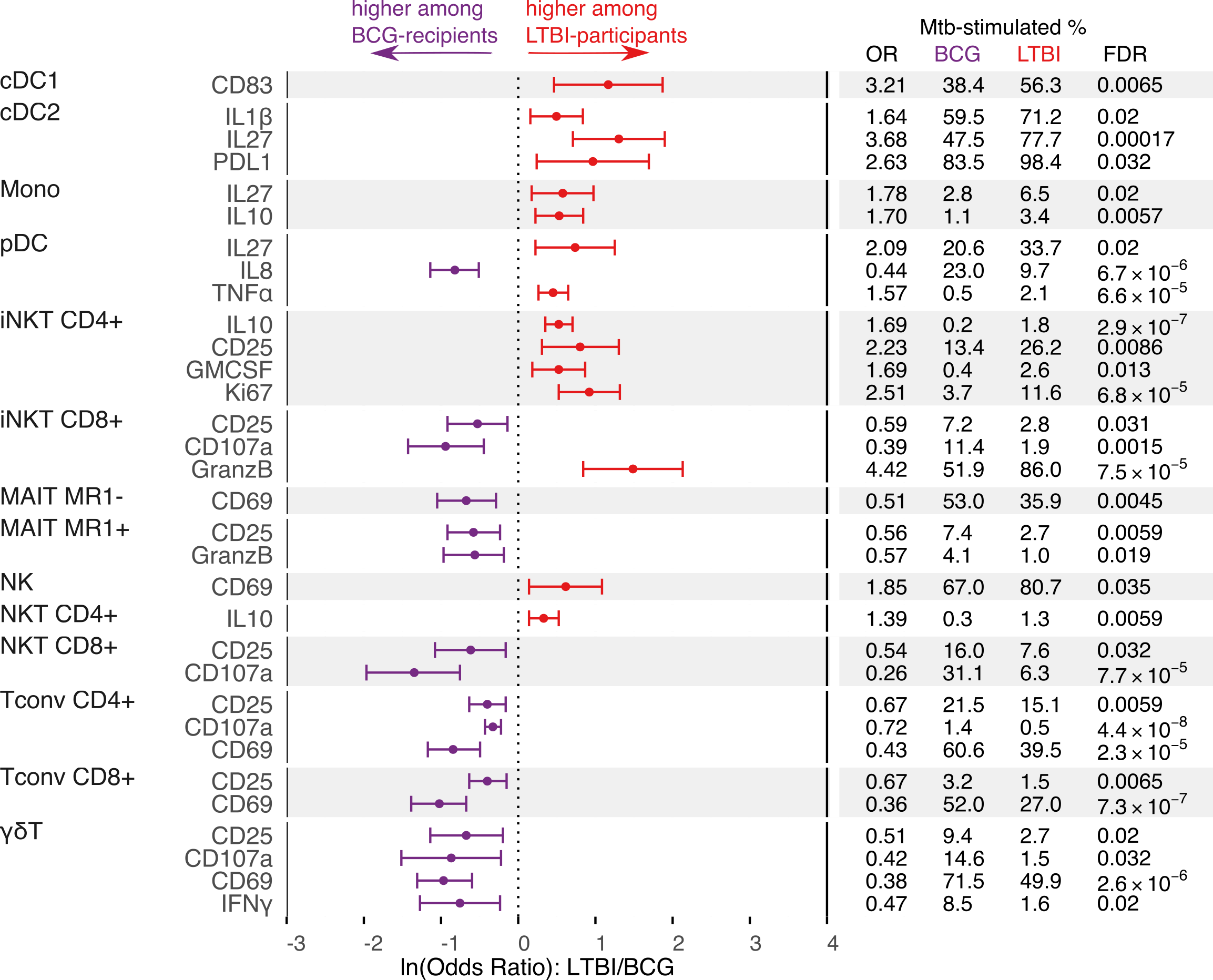

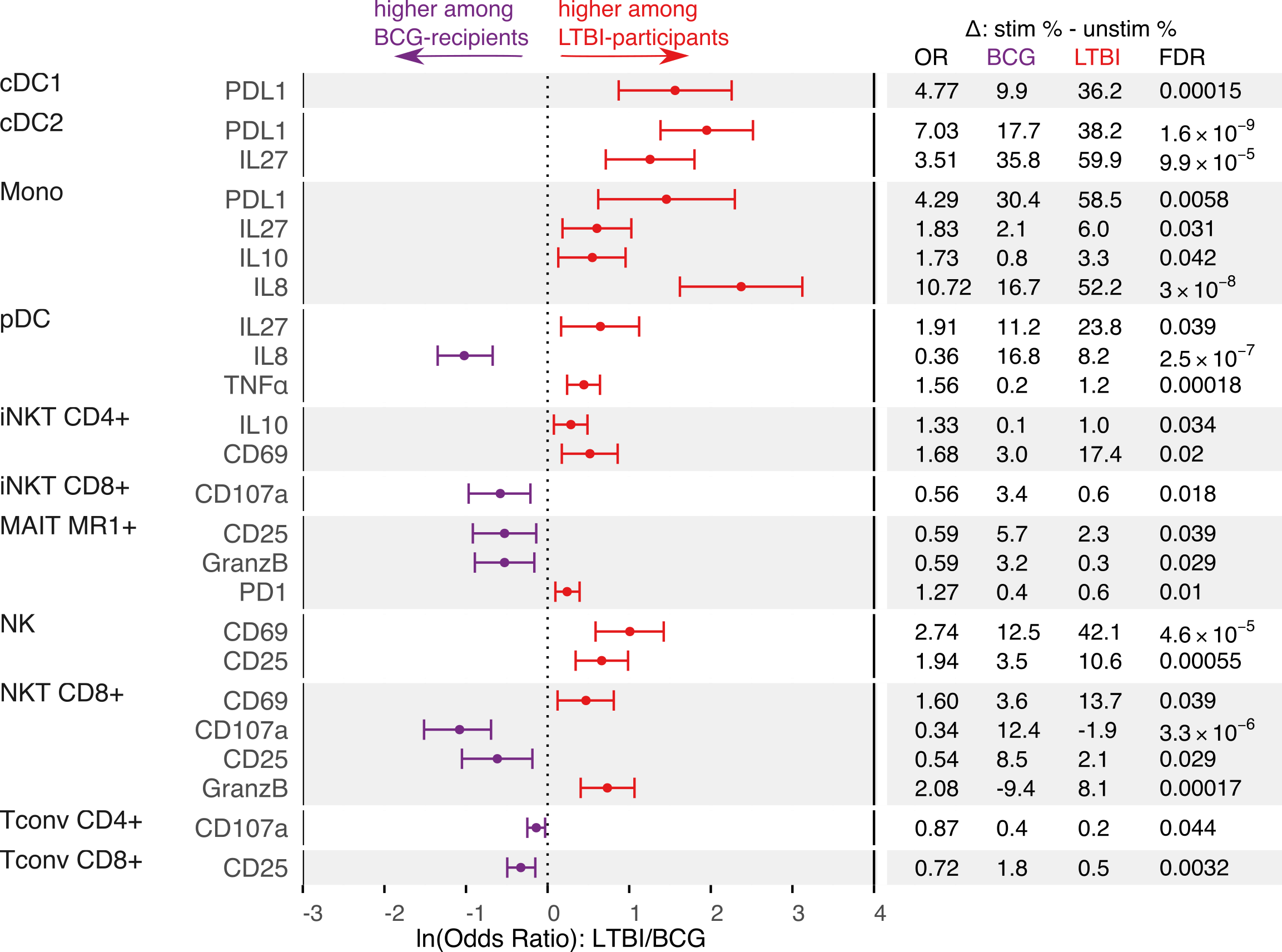

**Figure S5**. **Comparison of immune cell subset frequencies in BCG recipients and LTBI-articipants.** Data were derived from 11 LTBI-participants and 14 BCG-recipients. Forest plot displays ln-transformed odds ratios (lnOR) and 95% confidence intervals (CI). The dotted line shows no average effect (lnOR = 0, corresponding to OR = 1). Features on the right side of this line are more highly expressed among LTBI-participants (OR > 1), and features on the left side are more highly expressed among BCG recipients (OR < 1). The table shows the absolute OR and the means of each parameter in TB-resisters and LTBI-participants. **Top**: unstimulated PBMC; **Middle**: Mtb-stimulated; **Bottom**: Mtb-memory.

**A**

GD

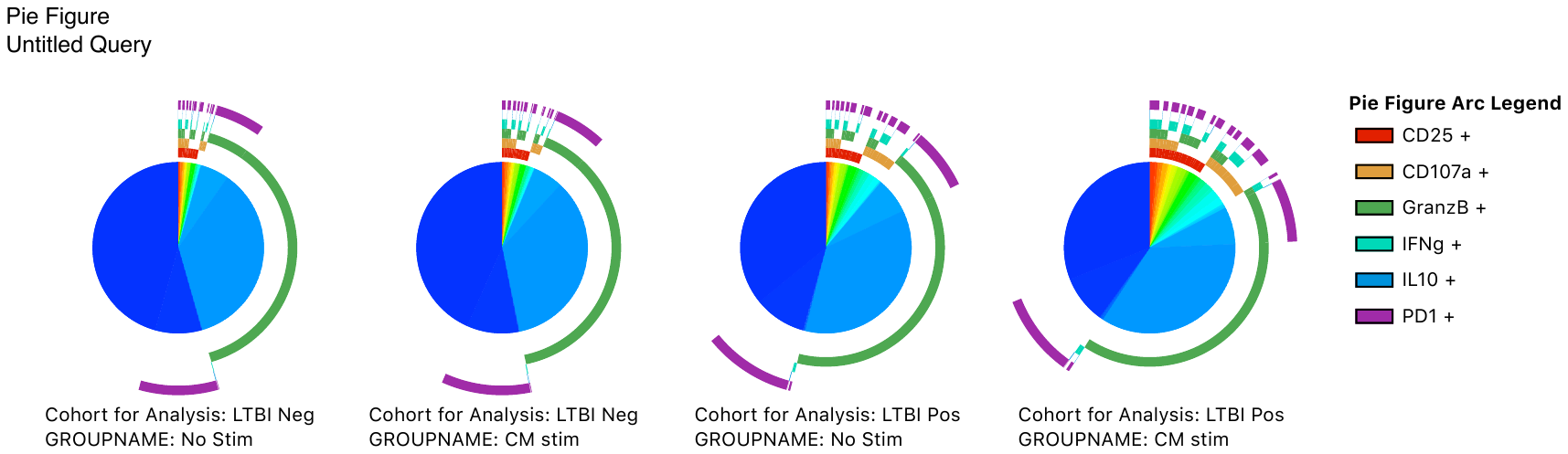

CD25

CD107a

GranzB

IFNγ

IL-10

PD-1

NK

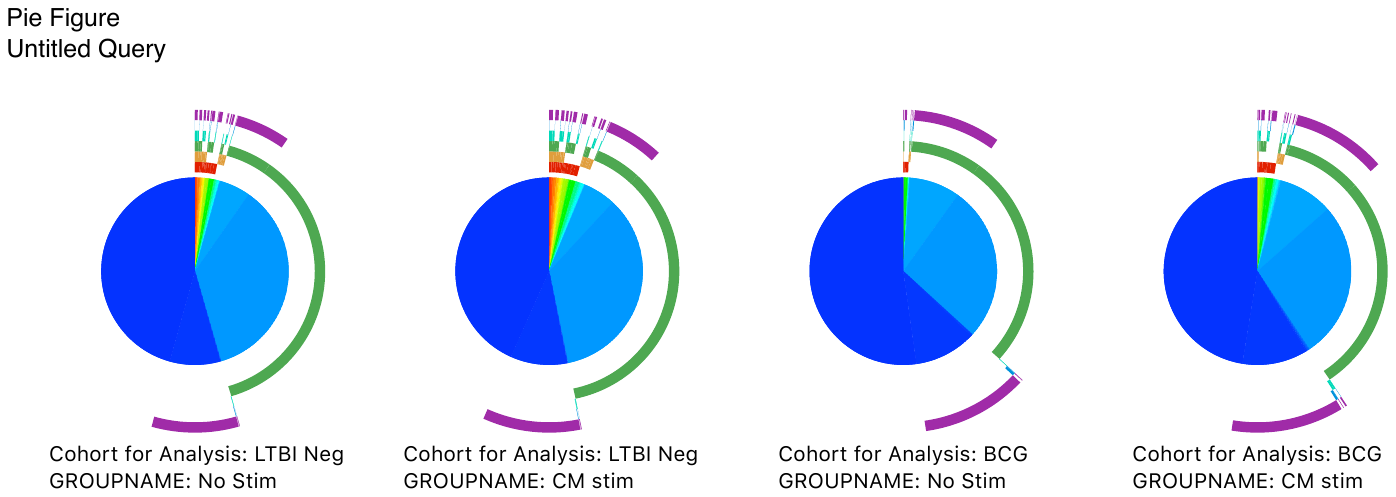

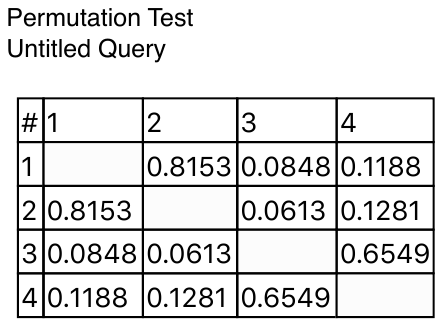

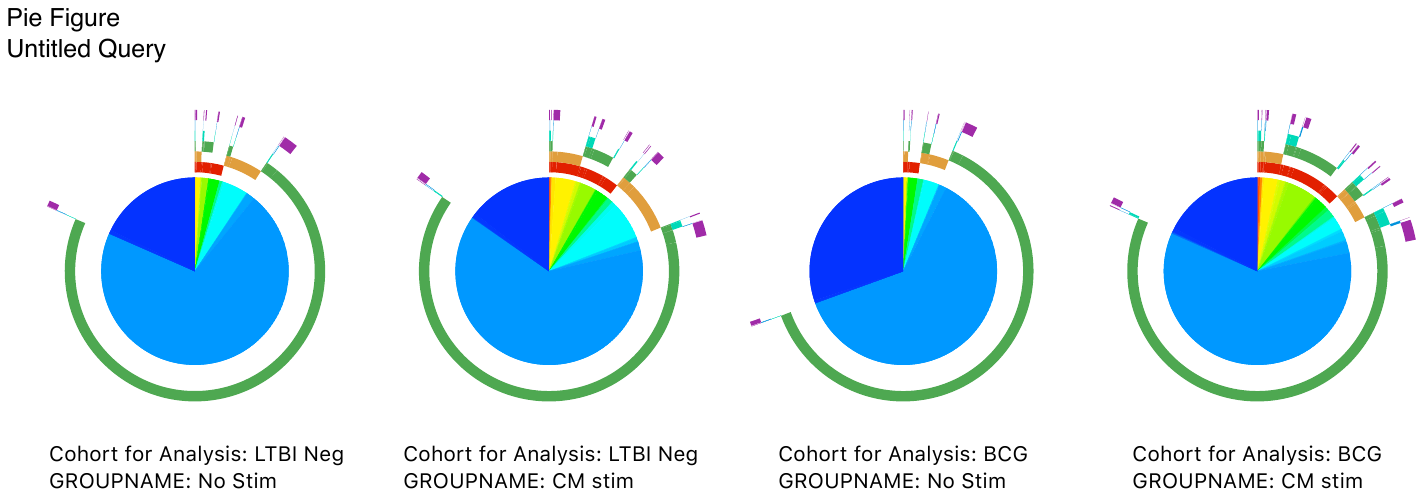

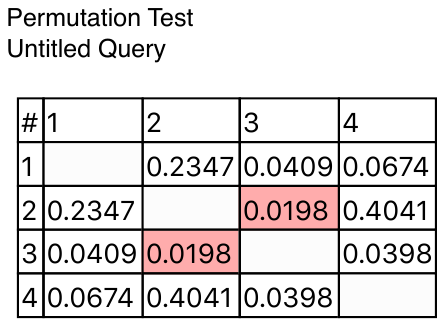

LTBIneg

NS

LTBIneg

CM

BCG

NS

BCG

CM

**B**

GD

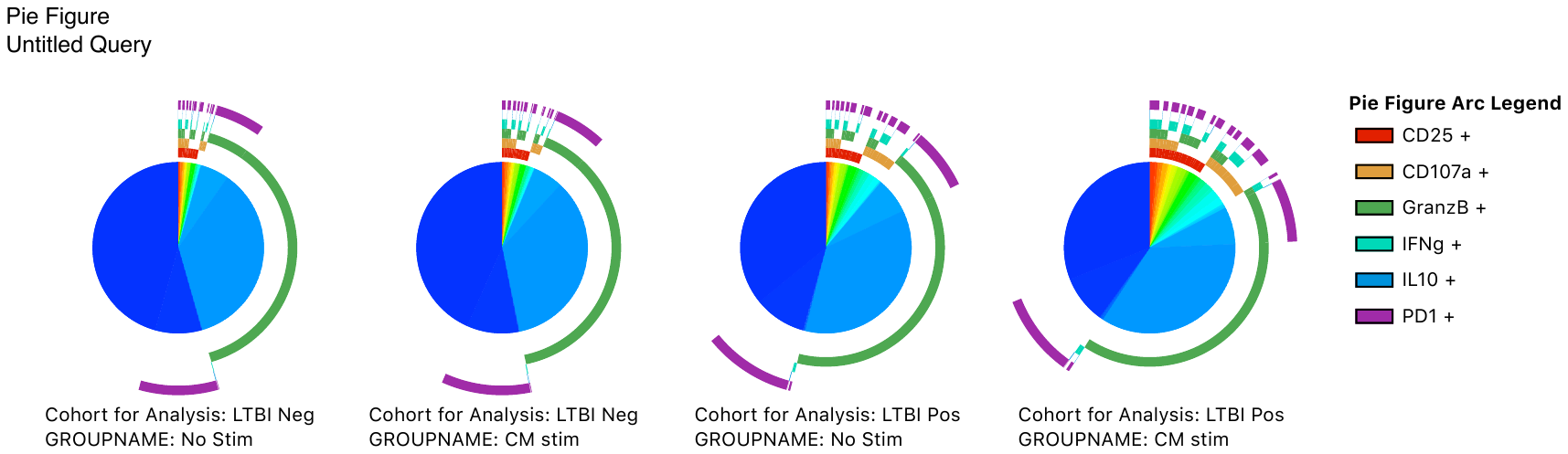

CD25

CD107a

GranzB

IFNγ

IL-10

PD-1

NK

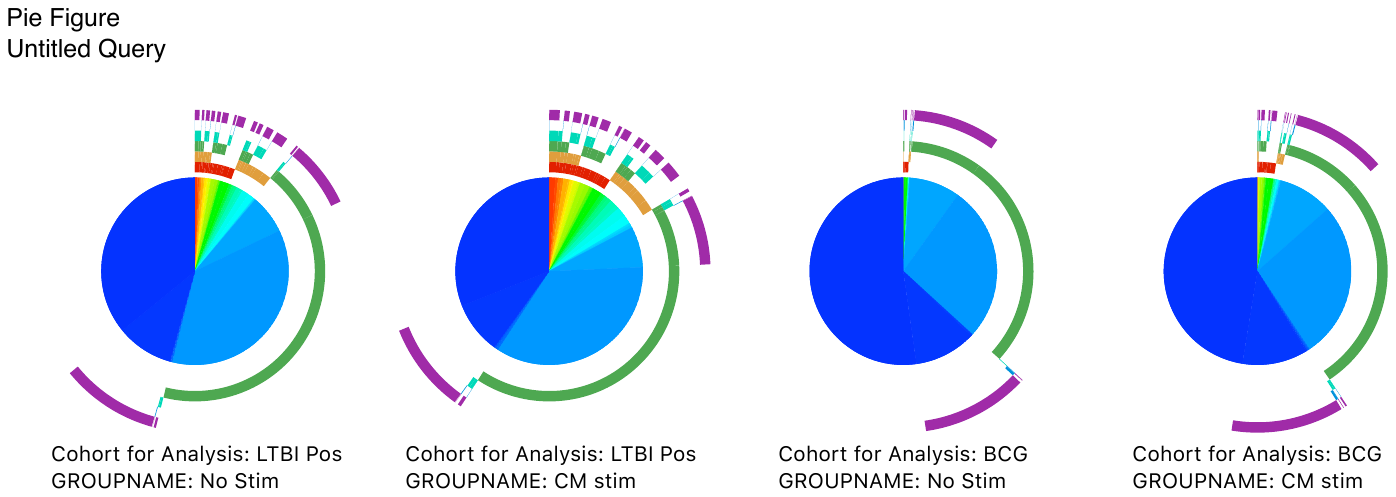

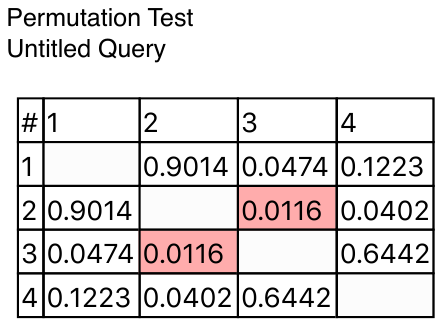

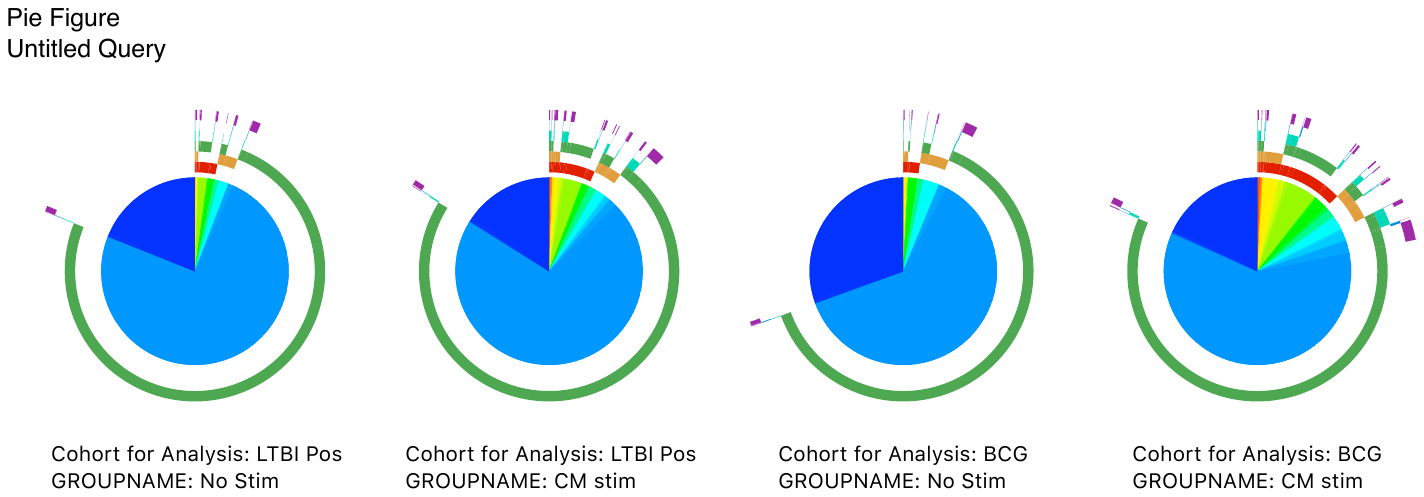

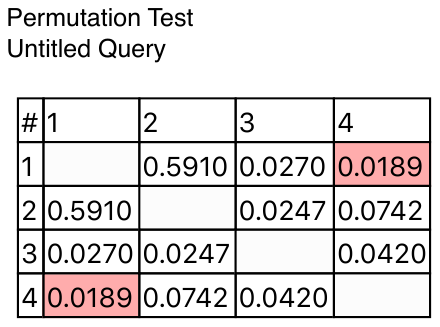

LTBIpos

NS

LTBIpos

CM

BCG

NS

BCG

CM

**Figure S6. Polyfunctionality of select T cell responses in BCG-recipients by comparison with TB-resisters (LTBIneg) and LTBI-participants (LTBIpos).** Data were derived from 14 BCG-recipients, 13 TB-resisters, and 11 LTBI-participants. **A:** BCG-recipients and TB-resisters; **B**: BCG-recipients and LTBI-participants. Pie charts 1-4 show proportions of cells expressing no markers (dark blue) or ≥1 marker (other colors) in response to Mtb stimulation or control. Arches show the markers expressed by the responding cells. The permutation tables present p values for comparisons across all groups and conditions. Tables show unadjusted p-values for each comparison. Pink boxes indicate significant differences defined by FDR p < 0.05.

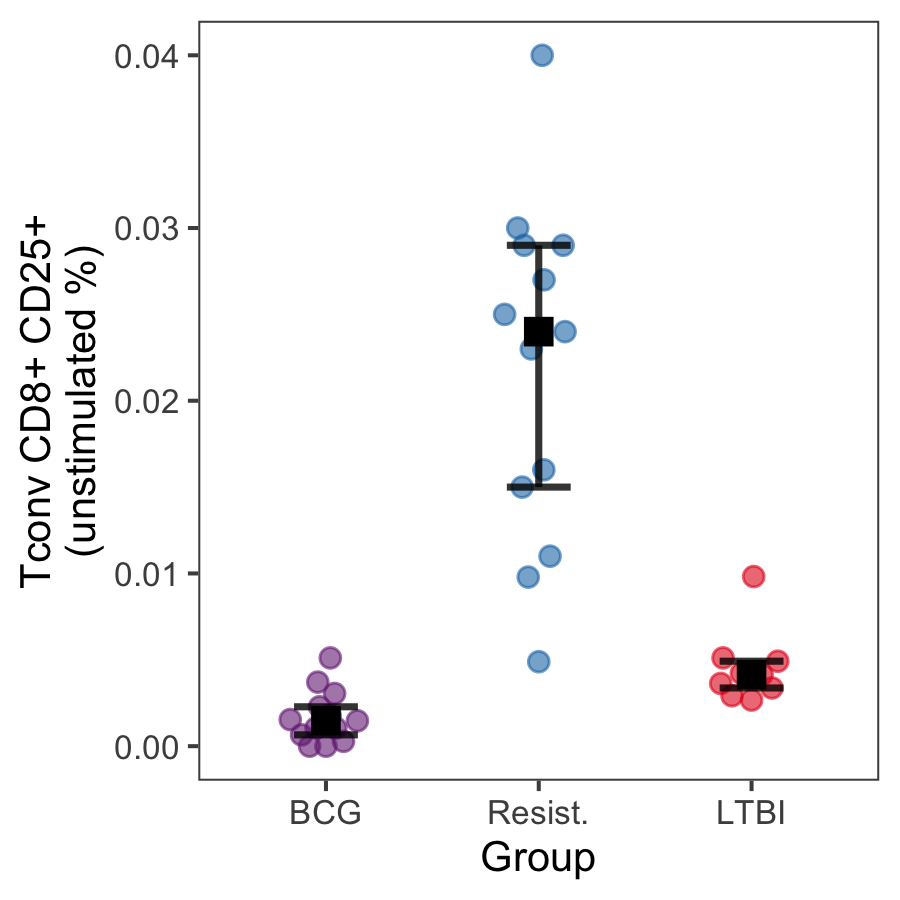

**Figure S7. CD8+GMM+GranzB+ T cells in BCG recipients compared with TB-resisters and LTBI-participants.** The graph shows results from 14 BCG-recipients, 13 TB-resisters and 9 LTBI-participants. Medians, upper and lower quartiles are indicated on the graph.
